# Lineage-Specific Epigenomic Mapping of Proximal and Distal Fallopian Tube Epithelium Reveals Mesenchymal/Wnt-Driven Tumor Phenotype Diversification

**DOI:** 10.1101/2025.08.27.672603

**Authors:** Liudan Wei, Huarong Wang, Wenshu Li, Jingyao Lian, Lijun Huang, Yifan Wang, Tianqi Li, Shiying Li, Zhenhua Hu, Zhaowei Tu, Wei Sun, Shuo Chen, Shuang Zhang

**Affiliations:** Department of Obstetrics and Gynecology, The Third Affiliated Hospital, Guangzhou Medical University, Guangzhou 510150, China; Guangdong Provincial Key Laboratory of Major Obstetric Diseases, The Third Affiliated Hospital, Guangzhou Medical University, Guangzhou 510150, China; Guangdong Provincial Clinical Research Center for Obstetrics and Gynecology, The Third Affiliated Hospital, Guangzhou Medical University, Guangzhou 510150, China; Guangdong-Hong Kong-Macao Great Bay Area Higher Education Joint Laboratory of Maternal-Fetal Medicine, The Third Affiliated Hospital of Guangzhou Medical University, Guangzhou 510150, China

**Author notes:** Corresponding author: Shuang Zhang; Shuo Chen; Wei Sun. Equal contribution (Liudan Wei, Huarong Wang).

## Abstract

High-grade serous ovarian cancer (HGSC) typically originates from the distal fallopian tube epithelium (disFTE), while the proximal epithelium (proFTE) remains less explored. Here, Using isogenic murine organoid models with common oncogenic drivers (*Tp53^−/−^; Kras^mut^; Pten^−/−^*), we show that disFTE and proFTE generate distinct tumor subtypes driven by region-specific transcriptional programs. Unlike disFTE, proFTE demonstrates continuous activation of epithelial-mesenchymal transition (EMT) across wild-type, mutant organoids, and derived tumors. Additionally, mutated proFTE organoids and their tumors exhibit persistent WNT signaling activation and grow independently of WNT signals. Chromatin profiling shows that mutant proFTE maintains a distinctive epigenetic profile with promoters associated with EMT and WNT pathways. Human proFTE with BRCAmut also shows epigenetic activation of EMT- and WNT-related genes. Tumors near the uterotubal junction (UTJ)—anatomically akin to proFTE—exhibit a pronounced EMT signature, increased Vimentin+E-cadherin+ and nuclear β-catenin expression. These findings suggest proFTE-derived tumors may occupy a unique niche in gynecological malignancies, with regional identity shaping tumor phenotypes.

## INTRODUCTION

Variations in histological and molecular characteristics across different zones of the same tissue play a crucial role in tumor development, incidence, and prognosis. Studies in cancers such as colon and prostate cancer have demonstrated that spatial heterogeneity within tissues can influence tumor initiation, progression, and therapeutic response^1–5^. Understanding these intratissue variations is critical for improving diagnostic accuracy and developing targeted therapeutic strategies. Anatomically, the fallopian tube (FT) is divided into two distinct parts: the distal segment adjacent to the ovary, comprising the infundibulum with associated fimbriae, ampulla, isthmus, and the proximal segment adjacent to the uterus, containing the isthmus and interstitial part ^6^. The distal fallopian tube (disFT), specifically fimbriae and ampulla region, are the focal point of research on high grade serous ovarian cancer (HGSC), which stands as the most prevalent and lethal form of ovarian cancer. This focus is driven by the frequent observation of serous intraepithelial carcinoma (STIC) in the secretory cells of disFTE in patients with Brca1/2 mutations, who are at a heightened risk of developing HGSC ^7–11^. STIC lesions are regarded as precursors to HGSC due to their shared morphological and molecular features, including the p53 signature and increased ki67 staining ^12–14^. Stem cell markers such as ALDH^hi^ and EPCAM^+^CD44^+^ITGA6^hi^Lin^-^, CD44^+^ peg cells are reported to be enriched in the distal fallopian tube, likely contributing to its vulnerability to oncogenic alterations and STIC formation ^15–17^. Although rare, there are reports indicating that lesions can also arise in the proximal FT (proFT) in BRCA1/2 carriers ^18^. Whether the proFT contributes to HGSC, and what are the molecular mechanisms distinguishing distal and proximal FT, especially when exposed to the same oncogenic alterations, remain unexplored.

Both distal and proximal part of FT, together with other urogenital system including uterus, and cervix, originates embryonically from the Müllerian duct ^19^. However, in adulthood, the distal and proximal FTE exhibit distinct cellular compositions and molecular profiles. The distal FT contains Pax8^+^ secretory cells and acetyl-α-tubulin^+^ ciliated cells, while the proFT predominantly expresses Pax8^+^ cells ^20^. In the disFT, Pax8^+^ secretory cells are broadly accepted as precursor cells that can differentiate into ciliated cells and transform under the influence of factors such as oncogenic incorporation, aging, and chemotherapy ^21, 22^. This transformation leads to secretory cell expansion and loss of ciliated cells, serving as biomarkers for early serous carcinogenesis within the fallopian tube ^23–25^. Despite the presence of ubiquitously Pax8^+^ cells in the proximal FT, their role during oncogenic transformation has largely been neglected. Notably, using lineage tracing on PAX2-GFP, in which GFP was observed only expressed in the proximal region of the oviduct, alongside with *PAX8^Cre/+^Rosa26^tdT/tdT^* lines, previous study indicate that distal and proximal FT represent intrinsically distinct lineages that follow separate differentiation trajectories and are maintained as independent entities ^26^. Whether the intrinsic lineage-specific differences between the distal FT and proximal FT contribute to variations in tumor-forming capabilities or properties, remain unknown.

Emerging single-cell ATAC-seq data from postmenopausal fallopian tubes have further highlighted significant lineage differences in chromatin structure and gene expression patterns between distal and proximal FTE ^27^. In patients with BRCA mutations, these epigenetic differences are exacerbated, with BRCA mutations inducing distinct regulatory changes in the two regions. Specifically, distal FTE undergoes rapid DNA methylation loss, particularly in HGSC-associated genes, whereas the proximal FTE exhibits different epigenetic changes that suggest unique regulatory mechanisms ^28^. These findings raise the intriguing possibility that BRCA mutations may drive region-specific tumorigenic processes, potentially influencing tumor subtype and pathological outcomes. However, the precise mechanisms underlying tumorigenesis in the proximal FT, as well as its ultimate pathological fate, remain unclear.

In this study, we aim to investigate the tumorigenic characteristics of epithelial cells in the proFTE and the disFTE, as well as the role of intrinsic lineage differences in this process. We generate organoid models derived from both distal and proximal FTE, introduce identical oncogenic alterations, and perform a comprehensive analysis of their tumor-forming potential, chromatin accessibility, and transcriptional programs. Through this approach, we uncover distinct characteristics of tumors originating from these two anatomically and molecularly distinct regions of the fallopian tube. In addition, we collected clinical human proFTE tumors and compared them with HGSC tumors and endometrial cancer (EC) tumors by immunostaining and single-cell RNAseq (scRNA-seq) analysis, which showed that the molecular characteristics of these tumors derived from different sites were significantly different. Our study provides insights into the lineage-specific mechanisms driving tumorigenesis and highlights how regional differences in epithelial cells contribute to tumor subtype and pathological outcomes.

## RESULTS

### Organoids Faithfully Recapitulate Lineage Differences in Distal and Proximal Regions of the Fallopian Tube

Investigating regional differences and their contribution to tumors in the same organ is difficult in vivo, we thus generated organoids derived from the fimbriae (disFTOs) and the isthmus (proFTOs) of the fallopian tube (Fig. 1a). Consistent with previous studies ^17, 26^, we observed that organoids derived from the distal region (disFTO) were significantly larger than those from the proximal region (proFTO) (Fig. 1b). Immunostaining for the two major cell types—secretory cells (Pax8^+^) and ciliated cells (acetyl-α-tubulin^+^)—in mouse fallopian tubes revealed that the distribution of ciliated cells gradually decreased from the distal to the proximal regions (Fig. 1c). Cross-sectional analysis showed a near absence of ciliated cells in the proximal isthmus and uterotubal sections. Correspondingly, disFTOs expressed both Pax8 and acetyl-α-tubulin, while proFTOs expressed ubiquitously Pax8 (Fig. 1c).

**Fig. 1.**
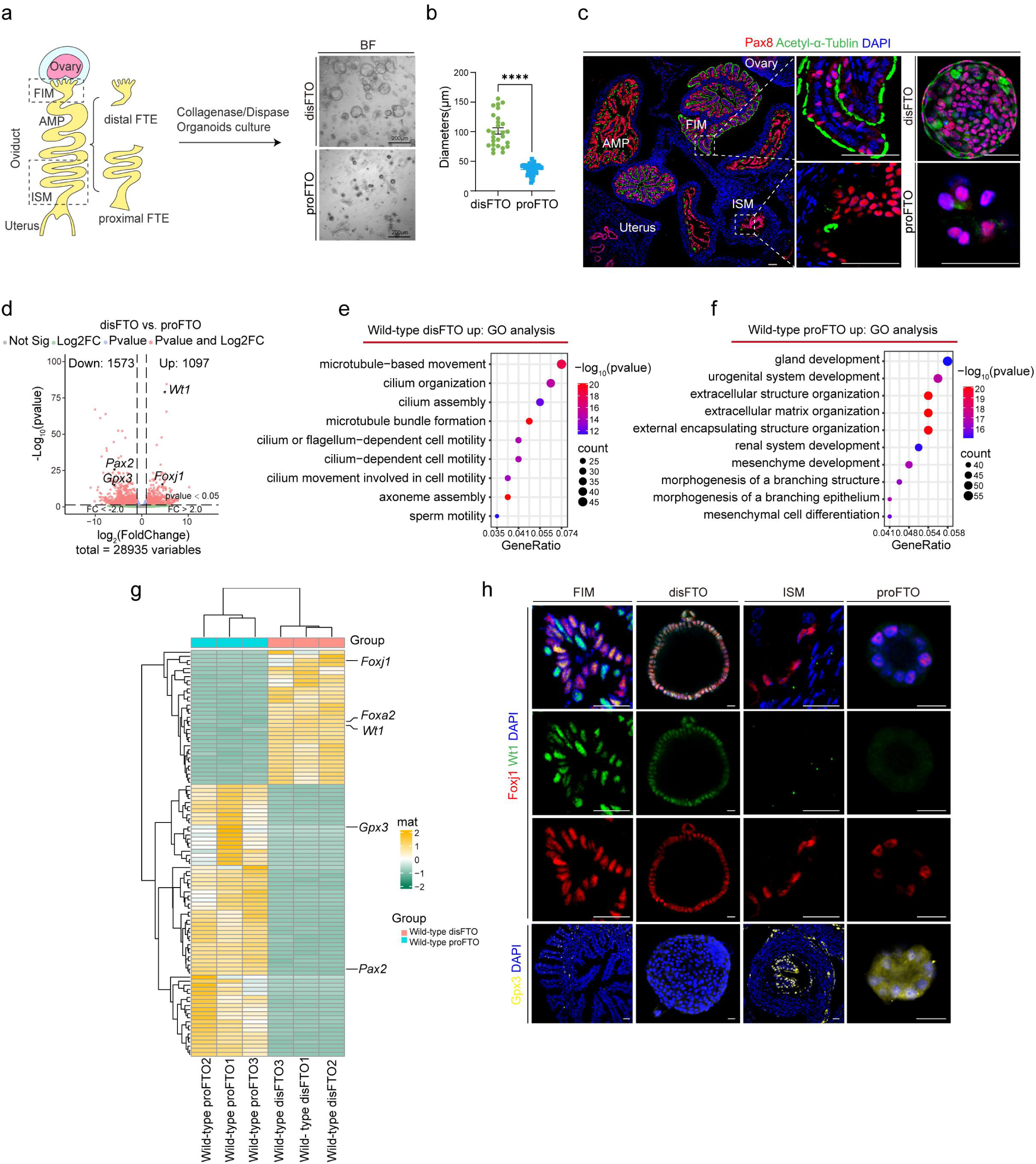
| Distinct Molecular and Morphological Characteristics of Distal and Proximal Fallopian Tube Organoids. **a** Schematic diagram illustrating the experimental design for deriving distal fallopian tube organoids (disFTO) and proximal fallopian tube organoids (proFTO) from mouse fallopian tubes. Bright-field (BF) images show disFTO and proFTO organoids cultured for 6 days. Scale bars represent 200 μm. FIM: fimbriae; AMP: Ampulla; ISM: Isthmus. **b** Quantification of the diameters of disFTO and proFTO organoids after 6 days of culture, data are presented as means ± SEM (unpaired t-test, ****P<0.0001). **c** Immunofluorescent staining of PAX8 and Acetylated α-tubulin (Acetyl-α-Tublin) on fallopian tube tissue and organoids from disFTO and proFTO cultured for 12 days. Scale bars represent 50 μm. **d** Volcano plot showing differentially expressed genes from RNA sequencing results comparing wild-type (WT) disFTO and proFTO after 6 days of culture. **e** Gene Ontology (GO) enrichment analysis of upregulated genes in WT disFTO compared to proFTO. **f** GO enrichment analysis of upregulated genes in WT proFTO compared to disFTO. **g** Heatmap displaying the top 100 differentially expressed genes between WT disFTO and proFTO based on p-values from RNA-seq results. The specific genes are shown in Table S4. **h** Immunofluorescent staining of the FIM, disFTO, ISM, and proFTO for markers Foxj1, Wt1, and Gpx3, with DAPI used for nuclear staining. Scale bars represent 20 μm.

To further investigate the molecular basis of these regional differences, we performed RNA sequencing on disFTOs and proFTOs derived from three mouse fallopian tubes (Fig. 1d). Among the 1,667 differentially expressed genes (DEGs) identified in disFTOs versus proFTOs (|LogFC| > 1, Pvalue < 0.05), 1097 were upregulated and 1573 were downregulated (Fig. 1d). In line with the morphological observations, Gene Ontology (GO) analysis revealed that disFTOs were enriched in terms related to microtubule-based movement, cilium organization, and cilium assembly—reflecting the differentiation of ciliated cells in the distal region (Fig. 1e). Conversely, proFTOs were enriched in terms associated with gland development, urogenital system development, and notably, mesenchymal development (Fig. 1f).

Consistent with previous reports, we identified the lineage-specific genes such as *Wt1*, *Foxj1*, *Gpx3*, and *Pax2*, were differentially expressed (Fig. 1g). Immunofluorescence staining further confirmed the expression of these genes in disFTEs, proFTEs, and their respective organoids (Fig. 1h). In addition, Wt1 exhibits exclusive expression in the distal fallopian tube across both human and murine models (Supplemental Fig. 1a, b). Together, these findings indicate that organoids derived from distal and proximal regions of the fallopian tube faithfully preserved the *in vivo* tissue-specific expression patterns and cellular composition, with disFTOs and proFTOs exhibiting distinct differences both morphologically and molecularly.

### Distinct Responses of disFTO and proFTO to Identical Oncogenic Integration

To determine whether disFTO and proFTO response differentially upon oncogenic cooperation, we generated organoid models from disFTE and proFTE using a PKC transgenic mouse model. This model harbors a *Tp53* knockout, an *LSL-Kras^G12D^* mutation, and *LSL-GFP-Cas9*, allowing for the expression of mutant *Kras* and *Cas9* upon Cre induction (Fig. 2a). In both disFTO and proFTO, GFP^+^ organoids were observed, indicating successful induction of *Kras* activation and *Cas9* expression by *Cre* incorporation (Fig. 2b). *Pten sgRNA* was then introduced, and single clones were selected. Western blotting and Sanger sequencing confirmed successful depletion of *Pten* in *Tp53^-/-^;Kras^G12D^ (*PK) disFTO and PK proFTO (Fig. 2c-f). *Kras* activation significantly increased the size of *Tp53^-/-^* organoids in both disFTO and proFTO, consistent with previous findings that *Kras* promotes proliferation ^29^. In contrast, *Pten* knockout significantly decreased organoid size and resulted in denser organoids in *Tp53^-/-^; Kras^G12D^; Pten^-/-^* (PKP) compared to PK organoids in both disFTO and proFTO (Fig. 2b, g, h).

**Fig. 2.**
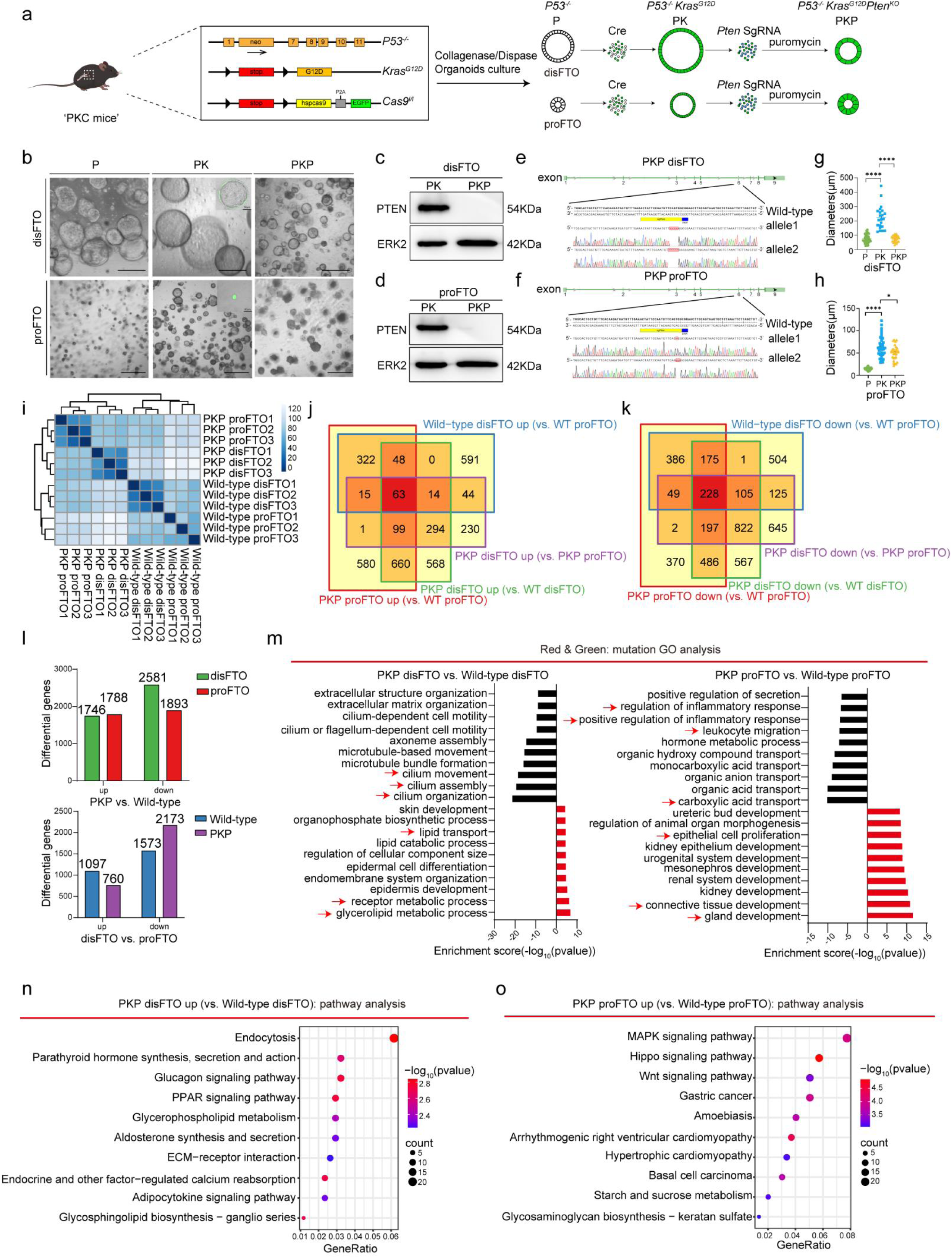
| Generation of Tumor Organoids Derived from Distal and Proximal Fallopian Tube Epithelium with Tp53, Kras, and Pten Alterations. **a** Schematic diagram illustrating the experimental design for generating distal fallopian tube organoids (disFTO) and proximal fallopian tube organoids (proFTO) with knockout mutations in Tp53 (P53^-/-^), Kras (Kras^G12D^), and Pten (Pten^-/-^), referred to as PKP organoids. **b** Bright-field images depicting disFTO and proFTO cultured for 6 days with indicated genotype. Scale bars represent 100 μm. **c-d** Western blot analysis confirming PTEN knockout in PKP disFTO **(c)** and PKP proFTO **(d)**. **e-f** DNA sequencing validation of Pten knockout in PKP disFTO **(e)** and PKP proFTO **(f)**. **g-h** Quantification of organoid diameters for disFTO **(g)** and proFTO **(h)** with various genetic alterations after 6 days of culture, presented as means ± SEM (unpaired t-test, ****P<0.0001, *P<0.05). **I.** Heatmap representing sample distances obtained via hierarchical clustering based on total gene expression levels across WT disFTO, WT proFTO, PKP disFTO, and PKP proFTO organoids. **j-k** Venn diagram illustrating the overlap of upregulated genes **(j)** and downregulated genes **(k)** between wild-type (WT) disFTO vs. WT proFTO, PKP disFTO vs. PKP proFTO, PKP disFTO vs. WT disFTO, and PKP proFTO vs. WT proFTO. **L** Upper panel: Differentially expressed genes impacted by PKP incorporation in both disFTO and proFTO. Lower panel: Comparative analysis of differential gene expression between disFTO and proFTO, with and without PKP alteration. The number of genes is indicated in each column of the charts. **m** GO enrichment analysis of differentially expressed genes in PKP disFTO compared with WT disFTO (left) and in PKP proFTO compared with WT proFTO (right). The black columns indicate pathways that are down-regulated, while the red columns represent pathways that are up-regulated. **n-o** Pathway analysis of upregulated genes in PKP disFTO relative to WT disFTO **(n)**, and in PKP proFTO relative to WT proFTO **(o)**.

We then performed RNA sequencing on PKP disFTO and PKP proFTO and compared these with their corresponding normal organoids. The heatmap showed clear separation between PKP FTOs and wild-type FTOs (Fig. 2i). Interestingly, oncogenic incorporation resulted in similar genetic changes in disFTO (1746 upregulated and 2581 downregulated genes) and proFTO (1788 upregulated and 1893 downregulated genes). However, the differences in gene expression between disFTO and proFTO (1573 downregulated and 1097 upregulated genes) were further exaggerated by oncogenic incorporation (760 downregulated and 2173 upregulated genes) (Fig. 2j-l).

Consistent with previous findings, oncogenic mutations inhibited the differentiation capacity of distal fallopian tube epithelial cells, as reflected by a significant downregulation of ciliated cell-associated genes and pathways ^23^. Immunofluorescence staining of Pax8 and acetyl-α-tubulin (a marker of ciliated cells) confirmed a marked reduction in ciliated cells in PK and PKP-disFTOs (Supplemental Fig. 2a, b). GO analysis further revealed that cilium-related processes, including cilium organization, cilium assembly, and cilium movement, were significantly downregulated in PKP disFTO compared to WT disFTO (Fig. 2m). In contrast, metabolic processes, including glycerolipid metabolism, receptor metabolism, and lipid transport, were enriched in PKP disFTO (Fig. 2m). Unlike mutant disFTO, oncogene-altered proFTO exhibited increased epithelium development, including gland development, connective tissue development, and epithelial cell proliferation, along with decreased carboxylic acid transport, leukocyte migration, and inflammatory response, indicating a more immunosuppressive environment in PKP proFTO (Fig. 2m). Furthermore, disFTO and proFTO with the identical oncogenic integration displayed divergent pathway enrichments. Mutant disFTOs were predominantly enriched in endocytosis, Glucagon signaling, and PPAR signaling pathways, whereas proFTOs showed significant enrichment in Hippo signaling, MAPK pathway, and Wnt signaling pathways (Fig. 2n, o).

### Mutant proFTOs Exhibit Wnt-Independent Growth and Mesenchymal features

Next, we directly compared the differentially expressed genes (DEGs) between PKP disFTO and PKP proFTO. As shown in Figure 3a, PKP disFTO exhibited a greater number of altered genes following oncogenic incorporation (2173 genes), whereas PKP proFTO only had 760 upregulated genes. Interestingly, PKP proFTOs were significantly enriched in mesenchymal development and Wnt signaling-related GO terms, as well as associated pathways, including Wnt, Hippo, and TGF-β signaling (Fig. 3b). Genes related to these pathways, such as *Cdh2*, *Zeb1*, and *Snai2*, up-regulated in PKP proFTO compared to PKP disFTO (Fig. 3c).

**Fig. 3.**
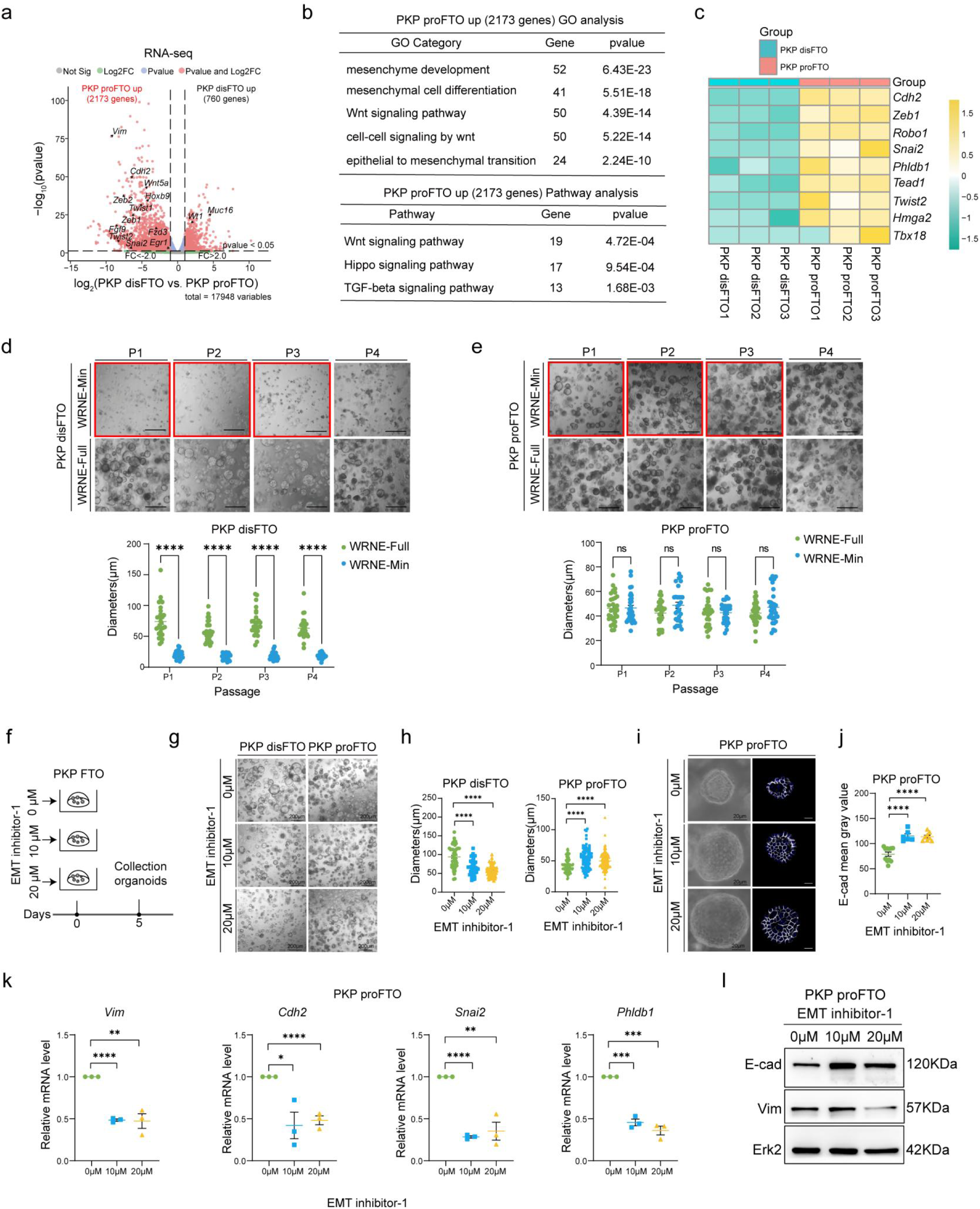
| The EMT and Wnt Signaling Pathways Are Significantly Enriched in PKP proFTO. **a** Volcano plot showing differentially expressed genes between PKP disFTO and PKP proFTO after six days of culture. **b** GO and pathway enrichment analyses of 2,173 genes upregulated in PKP proFTO compared with PKP disFTO. **c** Heatmap displaying the relative expression levels of genes associated with the Wnt signaling pathway. **d** Upper panel: Representative images showing the growth capacity of PKP disFTO in a Wnt-dependent manner. Organoids were passaged four times. Growth assay results represent three biological replicates. Scale bar: 200 μm. The red box highlights cells selected for the next passage. Lower panel: Quantification of organoid diameters for PKP disFTO cultured in WRNE-Full and WRNE-Min media. Diameters are presented as mean ± SEM (multiple unpaired t tests, ns: not significant, ****P < 0.0001). **e** Upper panel: Representative images showing the growth capacity of PKP proFTO in a Wnt-independent manner. Organoids were passaged four times. Growth assay results represent three biological replicates. Scale bar: 200 μm. The red box highlights cells selected for the next passage. Lower panel: Quantification of organoid diameters for PKP proFTO cultured in WRNE-Full and WRNE-Min media. Diameters are presented as mean ± SEM. ns: not significant. **f** Schematic diagram illustrating the experimental design of PKP FTO treated with EMT inhibitor-1 for five days. **g** Bright-field images of PKP disFTO and PKP proFTO treated with 10 μM and 20 μM EMT inhibitor-1 for five days. DMSO was used as the control. Scale bar: 200 μm. **h** Quantification of organoid diameters for PKP disFTO (left) and PKP proFTO (right) treated with EMT inhibitor-1 for five days. Data are presented as mean ± SEM. ****P < 0.0001. **i** Immunofluorescent staining of E-cadherin (E-cad) (white) in PKP proFTO treated with EMT inhibitor-1. DAPI (blue) was used for nuclear staining. Scale bar: 20 μm. **j** Quantitative analysis of E-cadherin fluorescence intensity in PKP proFTO (shown in **i**). Data are presented as mean ± SEM. ****P < 0.0001. **k** RT-qPCR analysis of gene expression levels relative to the EMT pathway in PKP proFTO. The scatter plot represents fold-change relative to EMT inhibitor-1 0μM, set as 1 (RQ, relative quantification). Error bars represent means ± SEM. β-actin was used as an endogenous control. n = 3 independent experiments, with triplicates for each gene per experiment. **P < 0.01; ***P < 0.001, ****P < 0.0001. **l** Western blot analysis of E-cadherin (E-cad), and Vimentin (Vim) expression in PKP proFTO treated with EMT inhibitor-1.

Given that Wnt activation is generally essential for organoid growth, and our culture system routinely includes Wnt3a, EGF, R-spondin-1, and Noggin (WRNE) to stimulate Wnt signaling. To determine whether reducing Wnt activation affects PKP proFTO growth, we compared organoid development under full WRNE medium (WRNE-Full) and Wnt-depleted conditions (WRNE-Min) in both PKP disFTOs and PKP proFTOs. Notably, PKP disFTO growth was significantly impaired in WRNE-Min, as indicated by a reduction in organoid size (Fig. 3d). However, this growth arrest was reversible upon reintroducing WRNE-Full medium. In contrast, PKP proFTOs showed no significant growth impairment under WRNE depletion (Fig. 3e), suggesting that mutated proFTOs are Wnt-independent. Furthermore, we applied EMT inhibitor-1 ^30^, which targets Hippo, TGF-β, and Wnt signaling pathways, to both PKP proFTOs and PKP disFTOs (Fig. 3f). As expected, EMT inhibitor-1 significantly reduced organoid growth in PKP disFTOs, as indicated by decreased organoid size (Fig. 3g, h), consistent with their Wnt-dependent nature. Interestingly, at the same dosage, PKP proFTOs exhibited distinct phenotypic changes, with increased organoid sizes and enhanced E-cadherin (E-cad) expression (Fig. 3g-j). Quantitative reverse transcription polymerase chain reaction (RT-qPCR) analysis of mesenchymal markers, including *Vimentin* (*Vim)*, *Cdh2*, *Snai2*, and *Phldb1*, further confirmed a reduction in mesenchymal properties in EMT inhibitor-treated PKP proFTOs (Fig. 3k). Western blot further verified the effect of an EMT inhibitor on PKP proFTOs (Fig. 3l), showing increased E-cad expression and decreased Vim levels in PKP proFTOs (Fig. 3l). These findings suggest that proFTOs possess inherent Wnt-independent and mesenchymal properties.

We then performed KEGG analysis using PKP disFTO up-regulated genes and revealed significant enrichment of pathways related to drug metabolism by other enzymes, and drug metabolism by cytochrome P450 in mutant disFTO (Supplemental Fig. 3a). These findings suggest that different transcriptional signatures between PKP disFTO and PKP proFTO may influence drug responses.

To explore the molecular basis underlying these differences, we analyzed key genes involved in drug resistance pathways (Supplemental Fig. 3b). The GST family was of particular interest, as it has been widely reported to be overexpressed in various cancers and implicated in chemotherapy resistance ^31, 32^. Consistently, our RT-qPCR analysis of GST family members, including *Gsto1*, *Gsto2*, *Gstt1*, and *Gsta1*, revealed significant upregulation in disFTO (Supplemental Fig. 3c).

To validate whether the higher expression of GST family affects drug responses, we conducted a drug screen using clinically relevant chemotherapeutic agents, including Gemcitabine, Carboplatin, Paclitaxel, Doxorubicin, and PARP inhibitors such as Niraparib, Rucaparib, and Olaparib, as well as platinums like Cisplatin (Supplemental Fig. 3d). Unexpectedly, these two lines showed equivalent IC_50_ values across most drug lines, except for Gemcitabine, where proFTO was more sensitive than disFTO (Supplemental Fig. 3d), indicating the drug sensitivity is influenced by not only GST family, but also by multiple factors.

### Proximal Tubal Tumors Exhibit Mesenchymal Traits and Wnt Activation Compared to Distal FT Tumors

Given the significant differences in gene expression and pathway enrichment between distal and proximal PKP-FTOs, we investigated whether tumor formation differs between these regions. Equal numbers of PKP-disFTO or PKP-proFTO cells were orthotopically injected into the mammary fat pads of nude mice (Fig. 4a). Luciferase imaging revealed that both PKP-disFTO-T and PKP-proFTO-T exhibited similarly robust tumorigenic capacity, with no significant difference in tumor formation efficiency (Fig. 4b, c). Strikingly, the two tumor types exhibited distinct morphological features: PKP-disFTO-T displayed serous-like characteristics, with fissured glandular structures infiltrating a fibrous stroma, whereas PKP-proFTO-T presented as solid cystic masses composed of scattered nests of mildly atypical epithelial cells embedded in dense fibrous tissue (Fig. 4d, Supplemental Fig. 4a). Despite their distinct morphological features, both tumor types uniformly expressed canonical HGSC markers (Pax8, CK7, Stathmin1, and Ki67), confirming their shared fallopian tube origin. Notably, WT1 - a marker typically associated with distal fallopian tube differentiation - was rarely detected in PKP-proFTO-T, highlighting their proximal origin-derived molecular characteristics (Fig. 4d, Supplemental Fig. 4a).

**Fig. 4.**
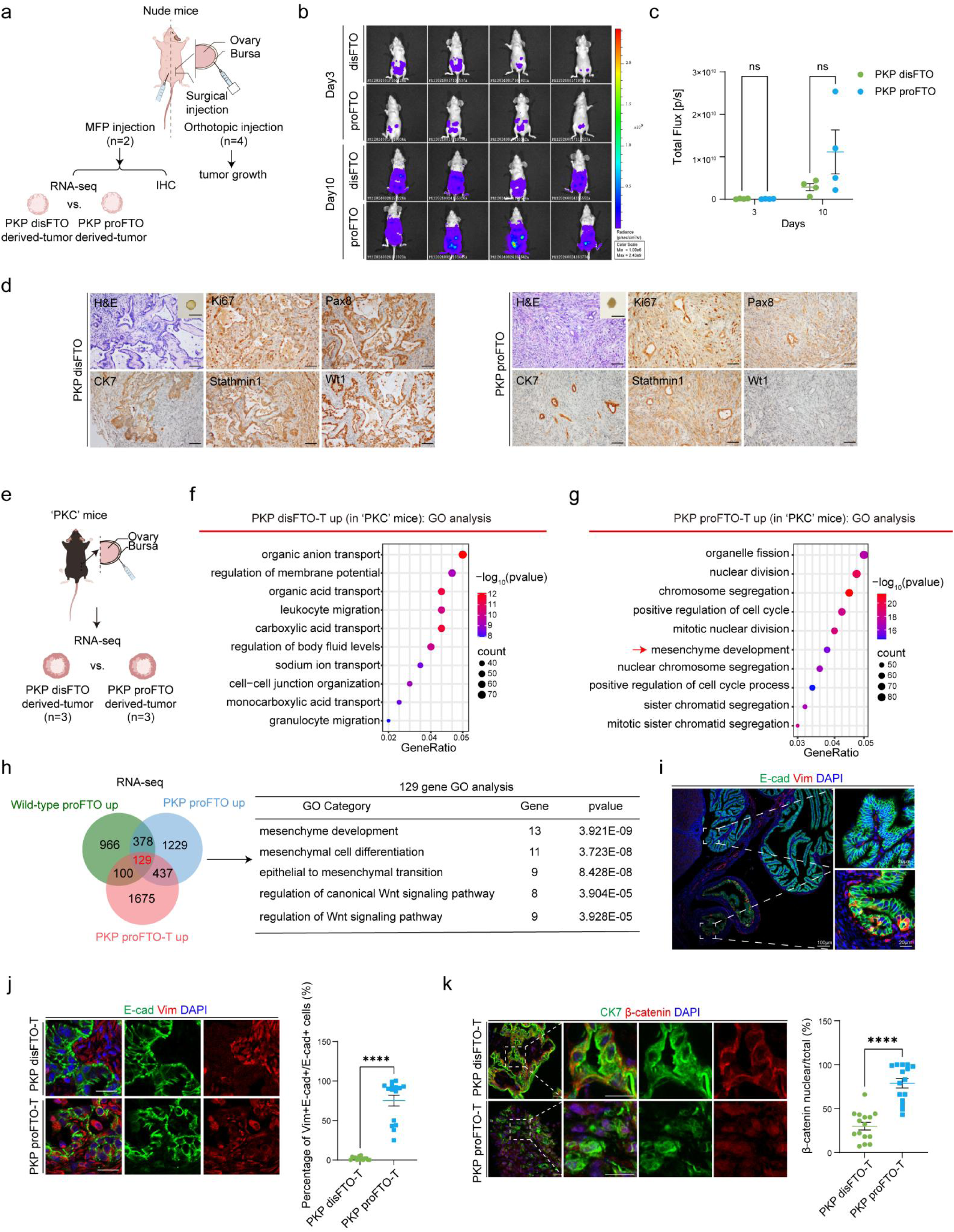
| Distinct Tumorigenic Characteristics of Distal and Proximal Fallopian Tube-Derived Organoids. **a** Schematic representation of orthotopic injection and mammary fat pad injection in 6-week-old nude mice. **b** Bioluminescence images of mice bearing orthotopic tumor allografts, measured by luciferase activity at 3 and 9 days after orthotopic injection (n=4). **c** Bioluminescence intensity of tumors in mice, 3 and 9 days post-orthotopic injection (n=4). Data are presented as mean ± SEM (multiple unpaired t tests, ns: not significant). **d** Representative images of organoid-derived tumors formed following mammary fat pad injection of PKP disFTO and PKP proFTO. H&E and IHC staining of tumors derived from PKP disFTO and PKP proFTO show the expression of markers associated with high-grade serous carcinoma (HGSC). Scale bars: 50 μm. The upper right panel of the H&E staining shows tumors formed by distal and proximal fallopian tube tumor organoids transplanted into mice. Scale bars: 1 cm. **e** Schematic representation of orthotopic injection in 6-week-old ‘PKC’ mice. Tumors were collected for RNA-seq analysis. **f** GO enrichment analysis of upregulated genes in PKP disFTO-T compared with PKP proFTO-T in ‘PKC’ mice. **g** GO enrichment analysis of upregulated genes in PKP proFTO-T compared with PKP disFTO-T in ‘PKC’ mice. **h** Venn diagram showing 129 commonly upregulated genes across three comparisons: wild-type proFT vs. wild-type disFT, PKP proFTO vs. PKP disFTO, and PKP proFTO-T vs. PKP disFTO-T. GO term and pathway enrichment analyses are shown for these overlapping genes. **i** Immunofluorescence staining of E-cadherin (E-cad), Vimentin (Vim), and DAPI in C57BL/6 mouse fallopian tube epithelium (FTE) (n = 3 tumors, count 5 fields of vision for each patient). Scale bars: 20 μm and 100 μm. **j** Immunofluorescence staining of tumors derived from PKP disFTO and PKP proFTO, labeled with E-cad, Vim, and DAPI. Scale bar: 20 μm. The percentage of E-cad and Vim double-positive cells among E-cad-positive cells in PKP disFTO and PKP proFTO tumor tissues is presented as mean ± SEM (unpaired t-test, ****P < 0.0001). **k** Immunofluorescence staining of tumors derived from PKP disFTO and PKP proFTO, labeled with CK7, β-catenin, and DAPI (n = 3 tumors, count 5 fields of vision for each patient). Scale bar: 20 μm. The percentage of nuclear β-catenin-positive cells among total β-catenin-positive cells in PKP disFTO and PKP proFTO tumor tissues is presented as mean ± SEM (unpaired t-test, ****P < 0.0001).

RNA sequencing of tumors from two mice per group revealed distinct molecular signatures. PKP-disFTO-T were enriched for pathways related to body fluid regulation and secretion, consistent with serous-like characteristics (Supplemental Fig. 4b). In contrast, PKP-proFTO-T showed enrichment for pathways including Notch and Wnt signaling and stem cell division, aligning with their in vitro signatures (Supplemental Fig. 4c).

To further investigate tumorigenic differences in the presence of a native tumor microenvironment, PKP-disFTO or PKP-proFTO were orthotopically injected into the ovarian bursa of ‘PKC’ mice (Fig. 4e). RNA sequencing of tumors from three mice per group identified 5,885 differentially expressed genes (LogFC > 1, P < 0.05) between PKP-disFTO-T and PKP-proFTO-T. GO analysis revealed that PKP-disFTO-T were enriched for leukocyte and granulocyte migration and body fluid regulation, suggesting a more immunoreactive microenvironment (Fig. 4f). In contrast, PKP-proFTO-T were enriched for pathways related to cell cycle regulation and mesenchymal development (Fig. 4g).

To identify conserved features of proximal fallopian tube-derived tumors, we integrated gene sets from wild-type proFTOs, PKP-mutated proFTOs, and their derived tumors, identifying 129 consistently upregulated genes. These genes were associated with mesenchymal development, epithelial-mesenchymal transition (EMT), and Wnt/Hippo signaling (Fig. 4h). IF staining for Vim, a mesenchymal marker, revealed its co-expression with E-cad in proFTE epithelial cells, a feature absents in distal FTE. This co-expression was more pronounced in PKP-proFTO-derived tumors, suggesting that proFTEs inherently undergo EMT, a process that persists following oncogenic transformation (Fig. 4i, 4k).

Additionally, we observed that β-catenin, a marker of active Wnt signaling, was rarely found in the nuclei of normal fallopian tube cells (Supplemental Fig. 5a). However, its presence significantly increased in tumors derived from PKP-proFTO compared to those from PKP-disFTO (Fig. 4k), suggesting that Wnt activation may play a crucial role in the tumor development of PKP-proFTO.

### Differential Chromatin Accessibility and Transcriptional Regulation between PKP-proFTOs and PKP-disFTOs

To uncover the regulatory mechanisms driving differential gene expression during tumorigenesis in the proximal and distal fallopian tubes, we analyzed chromatin accessibility and transcriptional activity using ATAC-seq and H3K27ac CUT&Tag-seq. ATAC-seq identified 9,584 chromatin loci with significantly higher accessibility in PKP-disFTO compared to PKP-proFTO, while only 2,285 loci were more accessible in PKP-proFTO (Fig. 5a). This aligns with prior studies in BRCA-mutate patients ^28^, where reduced DNA methylation was observed in the distal fallopian tube, suggesting distinct epigenetic modifications in the proximal and distal fallopian tube epithelium. Accessible chromatin regions were similarly distributed across regulatory elements in both regions, with promoter regions accounting for the largest proportion (Fig. 5b).

**Fig. 5.**
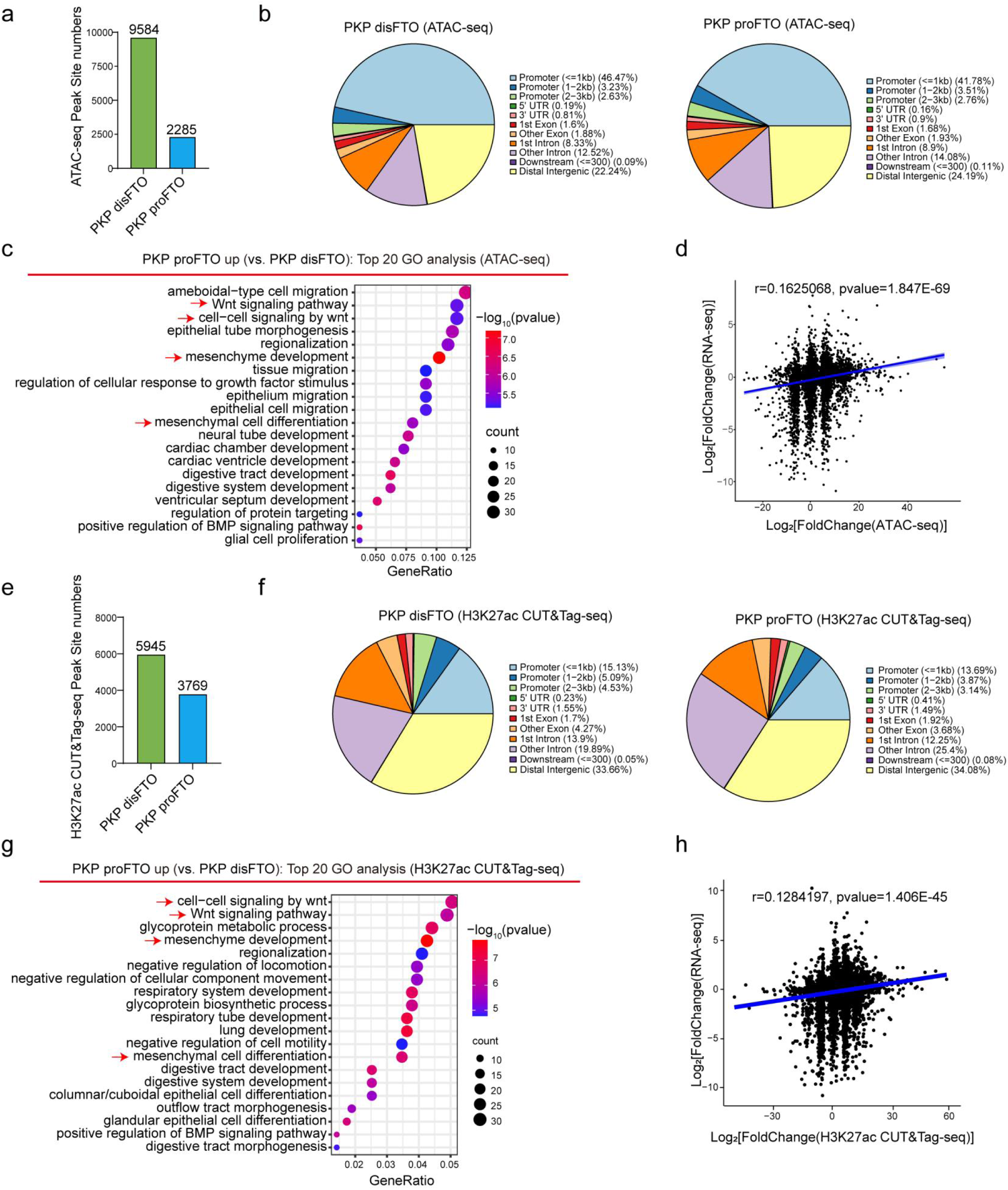
| The differential genes of PKP disFTO and PKP proFTO are regulated by chromatin Accessibility and transcriptional activity. **a** B*ar* graph displaying the number of specific ATAC-seq peak sites in PKP disFTO and PKP proFTO. **b** Location of PKP dFTO peak (left) and PKP pFTO peak (right) relative to the nearest annotated gene identified by ATAC-seq analysis. **c** Analysis of the top 20 GO-enriched genes up-regulated by PKP proFTO compared with PKP disFTO in the ATAC-seq promoter region. **d** Chromatin remodeling at gene promoters correlates significantly with changes in expression of the colocated genes (Pearson r = 0.1625068, pvalue = 1.847E-69). **e** B*ar* graph displaying the number of specific H3K27ac CUT&Tag-seq peak sites in PKP disFTO and PKP proFTO. **f** Location of PKP dFTO peak (left) and PKP pFTO peak (right) relative to the nearest annotated gene identified by H3K27ac CUT&Tag-seq analysis. **g** Analysis of the top 20 GO-enriched genes up-regulated by PKP proFTO compared with PKP disFTO in the H3K27ac CUT&Tag-seq promoter region. **h** H3K27ac CUT&Tag-seq at gene promoters correlates significantly with changes in expression of the colocated genes (Pearson r = 0.1284197, pvalue = 1.406E-45).

GO analysis of genes associated with differentially accessible promoter regions revealed enrichment of mesenchyme development and Wnt signaling pathways in the proximal fallopian tube, consistent with RNA-seq findings (Fig. 5c). H3K27ac CUT&Tag-seq further highlighted significant differences in transcriptionally active regions between the two regions (Fig. 5e, f). GO analysis of H3K27ac-modified promoter regions in proFTO confirmed enrichment of mesenchyme development and Wnt signaling pathways (Fig. 5g).

To determine whether differences in chromatin accessibility and H3K27ac modifications influence gene expression, correlation analysis showed a positive relationship between promoter chromatin accessibility, H3K27ac modification levels, and corresponding gene expression levels in both the proximal and distal fallopian tubes (Fig. 5d, h). These results suggest that oncogenic mutations drive region-specific gene expression patterns by modulating chromatin accessibility and H3K27ac-mediated transcriptional activation, with the mesenchymal and Wnt pathways being preferentially activated in the proximal fallopian tube.

### Epigenetic Regulation of PKP proFTO Reveals Enriched Wnt and Mesenchymal Transcription Factors and Genes

Building on the observed differences in chromatin accessibility and H3K27ac modifications, we further investigated how these epigenetic features regulate gene expression in proFTO. By integrating RNA-seq data with promoter-associated ATAC-seq and H3K27ac CUT&Tag-seq data, we identified 124 genes specifically upregulated in proFTO compared to disFTO (Fig. 6a). GO analysis of these genes revealed enrichment in pathways related to Wnt signaling, mesenchymal development, and regulation of Wnt signaling (Fig. 6a). Among these, 9 genes involved in mesenchymal development and 12 genes associated with Wnt signaling were consistently upregulated across all three datasets in proFTO (Fig. 6a, b).

**Fig. 6.**
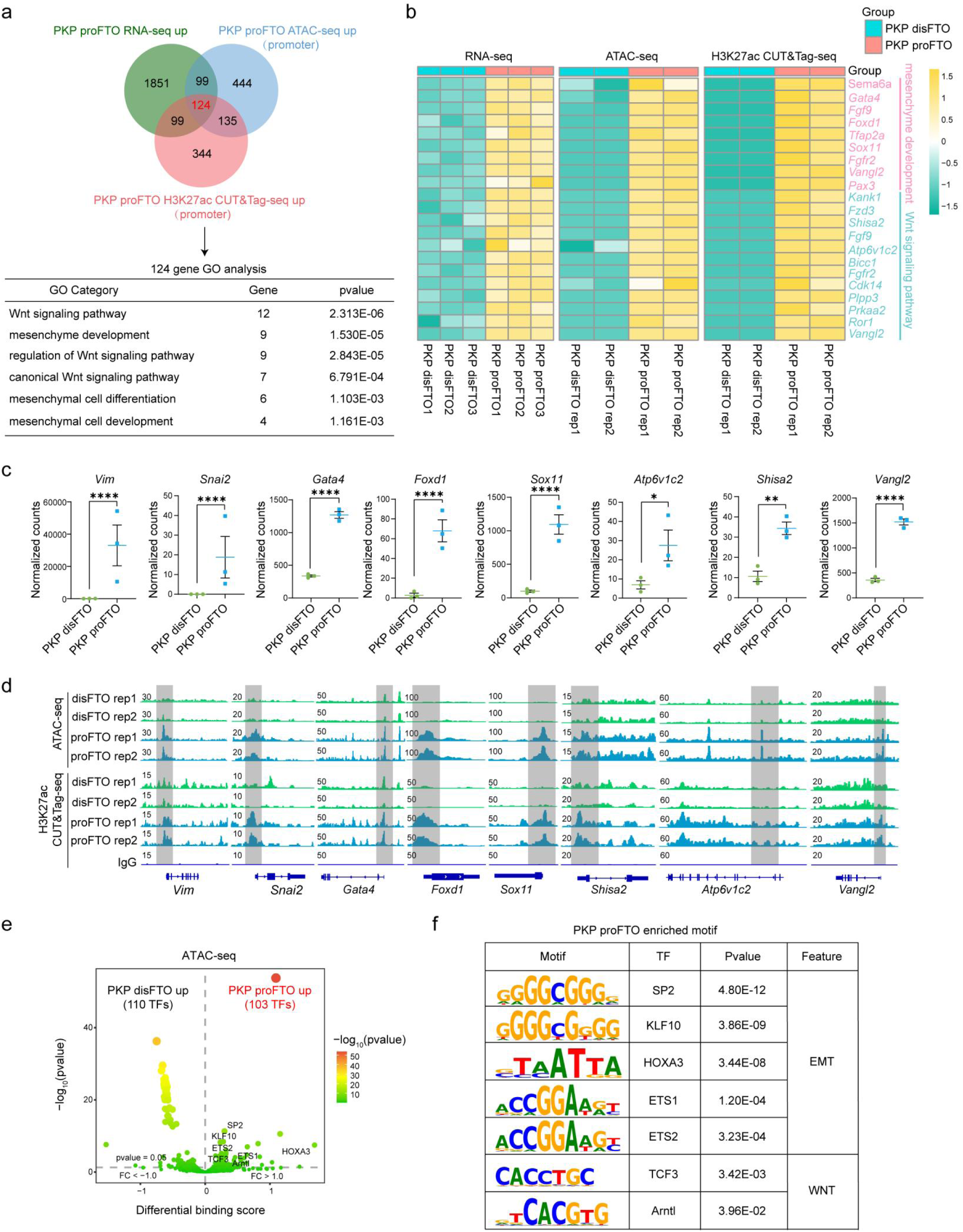
| Conservation of EMT and Wnt Signaling Pathway Activation in proFTO Validated by RNA-seq, ATAC-seq, and H3K27ac CUT&Tag-seq. **a** Venn diagram showing 124 up-regulated genes in PKP proFTO across three comparisons: RNA-seq, ATAC-seq in promoter, and H3K27ac CUT&Tag-seq in promoter. GO term and pathway enrichment analyses are shown for these overlapping genes. **b** Heatmap displaying the relative expression levels of 12 genes involved in both Wnt signaling pathway and 9 genes involved in mesenchyme development identified in figure 6A. **c** Normalized counts values of *Vim, Snai2, Gata4, Foxd1, Sox11, Shisa2, Atp6v1c2,* and *Vangl2* in PKP organoids. Relative differences in mRNA expression between distal and proximal fallopian tube organoids (unpaired t-test, *P < 0.05, **P < 0.01, ****P < 0.0001). **d** IGV visualization showing chromatin accessibility and transcriptional activity of *Vim, Snai2, Gata4, Foxd1, Sox11, Shisa2, Atp6v1c2,* and *Vangl2* in PKP proFTO, which are markedly higher than in PKP disFTO. **e** Volcano plot representing differentially enriched transcription factors (TFs) based on binding motif scores derived from ATAC-seq data. PKP proFTO shows enrichment of TFs associated with the EMT and Wnt pathways, including ETS1, ETS2, Arntl, HOXA3, TCF3, and the SP/KLF TF family. **f** Selected TF motifs specifically enriched in PKP proFTO, associated with the Wnt and EMT pathways.

To validate these findings, we examined normalized RNA-seq counts and visualized ATAC-seq and H3K27ac CUT&Tag-seq peaks for key genes implicated in mesenchyme development and Wnt signaling (e.g., *Vim, Snai2, Gata4, Foxd1, Sox11, Shisa2, Atp6v1c2,* and *Vangl2*) using the Integrative Genomics Viewer (IGV) (Fig. 6c, d). These analyses confirmed that increased chromatin accessibility and H3K27ac modifications in proFTO were associated with the upregulation of these genes, highlighting the role of epigenetic modifications in driving lineage-specific gene expression programs.

To explore whether these epigenetic differences influence transcription factor (TF) recruitment, we used TOBIAS software to analyze motif enrichment patterns specific to each region (Fig. 6e). Strikingly, 110 TF motifs were more enriched in disFTO, while 103 motifs were preferentially enriched in proFTO (Fig. 6e). Integrating motif enrichment data with RNA-seq revealed that TFs enriched in proFTO and associated with EMT-related genes included SP2, KLF10, HOXA3, ETS1, and ETS2 ^33–35^. Similarly, TFs linked to well established Wnt-related genes in proFTO included TCF3 ^36^ and Arntl ^37^ (Fig. 6e, f). These findings suggest that differential chromatin accessibility and H3K27ac modifications between proximal and distal fallopian tubes selectively recruit distinct TFs, thereby driving the specific activation of EMT- and Wnt-related gene expression programs in proFTO.

### Regional Identity of proFTE Drives Molecular Divergence from Typical HGSC and EC

To explore potential differences in epigenetic regulation between the distal and proximal regions of the human fallopian tubes, we analyzed data from a previous study ^28^ and conducted a GO analysis on hypomethylated genes, which are generally associated with transcriptional activation, at the TSS in the proximal tube compared to the fimbrial (distal) tube (Fig. 7a). The GO analysis revealed significant enrichment in biological processes such as mesenchymal development, morphogenesis, cell differentiation, EMT, and Wnt signaling pathways (Fig. 7a). These findings align with our earlier observations in mice (Fig. 4h-k; Fig. 5c, g).

**Fig. 7.**
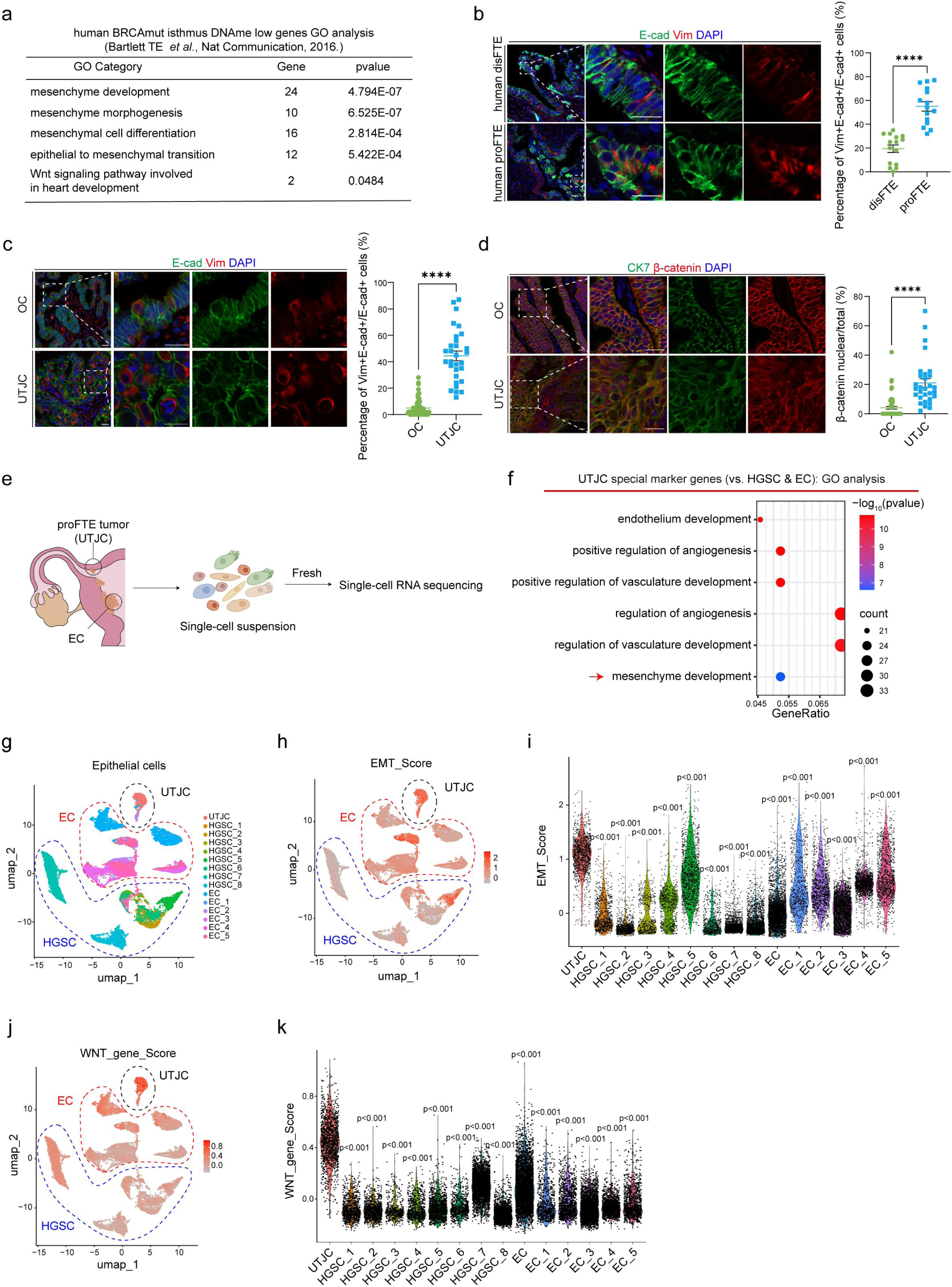
| Human Proximal Fallopian Tube Tumors Manifest Distinct Molecular Signatures Divergent from EC and OC. **a** Using data from Bartlett TE *et al*., 2016 published in nature communication. GO analysis of data in the transcription start site (TSS) region using Top CpGs, which are hypermethylated in fimbrial compared to proximal tube in BRCA1/2 mutation carriers. **b** Immunofluorescence staining of human distal fallopian tube epithelium (disFTE) and proximal fallopian tube epithelium (proFTE) with E-cad, Vim, and DAPI. Scale bar: 20 μm. The percentage of E-cad and Vim double-positive cells among E-cad-positive cells in human disFTE and proFTE (n = 3 patient samples, count 5 fields of vision for each patient) is presented as mean ± SEM (unpaired t-test, ****P < 0.0001). **c** Immunofluorescence staining of OC, and UTJC with E-cad, Vim, and DAPI (n = 4 – 15 patient samples, count 5 fields of vision for each patient). Scale bar: 20 μm. The percentage of E-cad and Vim double-positive cells among E-cad-positive cells in human OC, and UTJC is presented as mean ± SEM (unpaired t-test, ns: not significant; ****P < 0.0001). **d** Immunofluorescence staining of OC, and UTJC with CK7, β-catenin, and DAPI (n = 4– 15 patient samples, count 5 fields of vision for each patient). Scale bar: 20 μm. The percentage of nuclear β-catenin-positive cells among total β-catenin-positive cells in human OC, and UTJC is presented as mean ± SEM (unpaired t-test, ****P < 0.0001). **e** Workflow depicting the collection and processing of specimens of human proFTE tumor (UTJC), and EC for scRNA-seq. **f** GO enrichment analysis of special marker genes in UTJC compared to HGSC and EC. **g** The UMAP plot demonstrates the epithelial cells in UTJC, HGSC, and EC. **h** Scatterplots show the EMT transcriptomic characteristics of epithelial cells in UTJC, HGSC, and EC. The analytical method of EMT score was referenced from previous report ^40^. **i** Violin plots quantify EMT scores for UTJC, HGSC, and EC. **j** Scatterplots show the WNT transcriptomic characteristics of epithelial cells in UTJC, HGSC, and EC. **k** Violin plots quantify WNT gene scores for UTJC, HGSC, and EC.The EMT and WNT score genes are shown in table S5.

To further confirm this, we collected tissue samples from the human fallopian tube, specifically from the distal fimbriae (disFT) and the proximal isthmus near the uterine-tubal junction cancer (UTJC, classified as tumors located at the uterine-tubal junction). We also obtained samples from ovarian cancer (OC), UTJC, and endometrial cancer (EC). Using immunofluorescence staining for E-cad and Vim, markers of EMT, we identified regional differences in the presence of E-cad^+^Vim^+^ double-positive cells. Notably, the proFTE exhibited a significantly higher proportion of these double-positive cells (Fig. 7b), mirroring patterns observed in mouse models.

Next, we examined whether the abundance of E-cad^+^Vim^+^ cells correlate with tumor origin. Remarkably, these double-positive cells were most abundant in UTJC, which corresponds to the proFT. In contrast, they were least abundant in OC, derived from the disFT, and showed intermediate levels in EC, originating from the endometrium (Fig. 7c, Supplemental Fig. 6a). These patterns are consistent with the relative expression profiles observed in normal human disFTE and proFTE (Fig. 7b).

Consistent with our mouse data, nuclear β-catenin translocation was rare in both proximal and distal normal human fallopian tubes (Supplemental Fig. 5b). However, UTJC exhibited higher nuclear β-catenin translocation compared to OC, although this difference was not statistically significant (Fig. 7d). Notably, EC showed significantly higher β-catenin nuclear translocation compared to UTJC (Supplemental Fig. 6b). These findings suggest that Wnt signaling becomes distinctly activated in tumors originating from the proximal fallopian tube, paralleling observations in mouse models.

Clinically, we identified a patient presenting with both a uterine-tubal junction tumor (UTJC) and disseminated endometrial carcinoma, with the UTJC located within the proximal fallopian tube region. It remained unclear whether the UTJC originated from metastatic spread of endometrial cancer or arose independently from the proximal fallopian tube epithelium (proFTE). To investigate this, we performed cytoreductive surgery with curative intent to resect both the endometrial and proFTE-associated lesions (Fig. 7e). Tumor samples from these sites underwent single-cell RNA sequencing (scRNA-seq). For comparative purposes, we integrated these datasets with publicly available scRNA-seq profiles from HGSCs (n = 8) and ECs (n = 5) ^38, 39^.

Unsupervised clustering of the combined dataset revealed that UTJC tumor cells formed a distinct cluster, clearly separable from both HGSC and EC (Fig. 7g). GO analysis of UTJC-specific genes highlighted the enrichment of pathways associated with angiogenesis and mesenchymal development (Fig. 7f). Furthermore, quantitative analysis of EMT scores and WNT relative gene scores confirmed that UTJC tumor cells exhibited significantly higher EMT and WNT activity than those of HGSC and EC (Fig. 7h-k) ^40^, paralleling observations in results of mouse model and immunostaining (Fig. 7b, c). Collectively, these findings suggest that proFTE-derived tumors may represent a distinct biological niche within the spectrum of gynecologic malignancies, where regional anatomical identity significantly shapes their molecular and phenotypic characteristics.

## DISCUSSION

Differences in histology and molecular profiling across distinct zones of the same tissue underscore their potential contributions to tumor development, incidence, and prognosis ^1–5^. While previous research has documented anatomical and cellular heterogeneity along the distal-proximal axis of the fallopian tube ^17, 26^, the implications of these variations for tumor formation, characteristics, and treatment response remain underexplored. Our study demonstrated that proFTOs exhibited distinct biological traits—such as Wnt-independent growth and mesenchymal characteristics—compared to their distal counterparts. Furthermore, tumors derived from PKP-deficient proFTOs displayed increased Wnt activation and pronounced mesenchymal signatures, suggesting that anatomical origin may influence phenotypic diversity in fallopian tube-derived malignancies. However, it remained unclear whether these differences translated into clinically significant variations in tumor behavior or treatment responses. Future studies that correlate anatomical origin with patient outcomes and therapeutic vulnerabilities will be crucial in determining if proximal-derived tumors warrant classification as a distinct entity or if they represent part of the broader spectrum of fallopian tube-originating malignancies (Fig. 8).

**Fig. 8.**
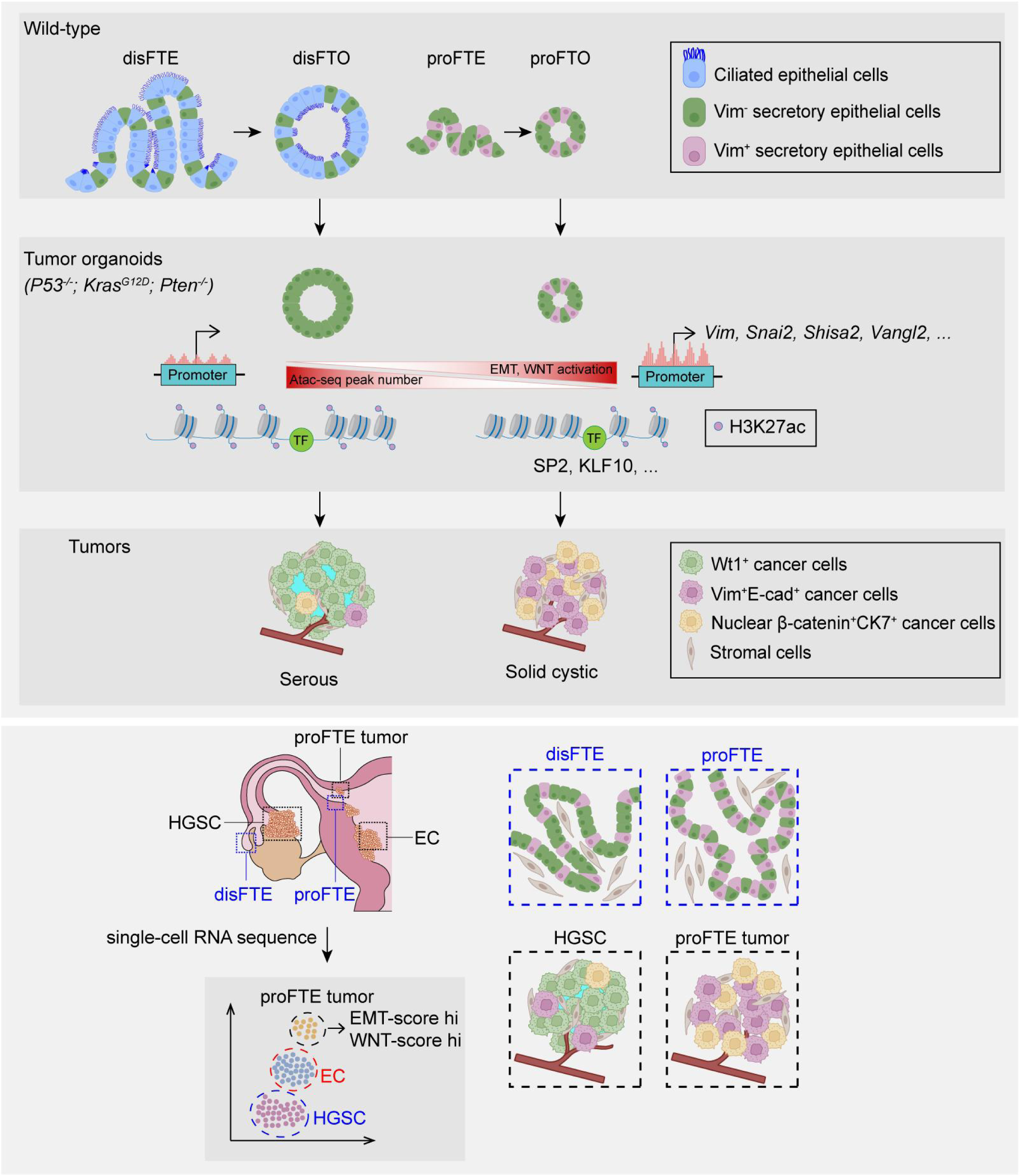
| Graphical summary of the differences between the distal and proximal fallopian tubes in human and mice.

Consistent with the widely accepted notion that the disFT is the cell of origin for HGSC, our study found that PKP-disFTO-derived tumors are enriched for pathways related to body fluid regulation and secretion, as well as enriched in genes associated with immunoglobulin secretion, lipid metabolism, cytokine production, and inflammatory responses. This aligns with the disFT’s location in the lipid-rich peritoneal cavity and its exposure to cytokine- and chemokine-rich follicular fluid during ovulation ^16, 41, 42^. In contrast, the proFT is relatively anoxic and enriched in mesenchymal development and EMT-related genes, such as *Vim,* and *Snai2*, as well as pathways like Hippo signaling. Intriguingly, treatment with EMT inhibitor-1 induced a mesenchymal-to-epithelial transition (MET) in PKP proFTOs, as evidenced by increased E-cadherin expression and decreased mesenchymal markers, indicating that mesenchymal features play a crucial role in proFTO identity. Remarkably, prior studies have shown that disFT is less methylated than proFT in BRCA mutation carriers ^28^, which aligns with our findings that PKP-disFTOs exhibit greater chromatin openness and transcriptional activity compared to PKP-proFTOs. This suggests that disFT has higher transcriptional activity than proFT during cancer evolution, and the higher transcriptional activity could contribute to the aggressive nature of HGSC.

A previous study demonstrated that mesenchymal reporter-positive epithelial cells are stably maintained in the oviduct epithelium of adult mice across multiple pregnancies ^43^. Similarly, through re-clustering of FTE cells from healthy donors, a recent work identified a non-ciliated secretory cell subpopulation co-expressing both epithelial and mesenchymal markers ^20^. In this study, we showed higher mesenchyme development and EMT activity in proFTE compared to disFTE in both murine and human samples, the co-expression of epithelial and mesenchymal markers was more pronounced in mutated proFTOs and their derived tumors, indicating the EMT state in the proFT is intrinsic and stable.

We revealed that tumors originating from the proFT molecularly and clinically resembled EC. Several factors contribute to this resemblance. Anatomically, the proximal fallopian tube is located adjacent to the uterus, serving as a conduit between the two structures. Furthermore, proFTOs display activated Wnt signaling and reduced dependence on Wnt factors, a characteristic also observed in endometrial organoid cultures ^45, 46^. Pathologically, a significant percentage of EC patients present with abnormalities in the proFT ^47, 48^. Notably, while UTJC and EC shared overlapping genetic features—often leading to diagnostic overlap—key molecular distinctions exist. For instance, ProFT-derived tumors displayed higher EMT scores, wheras EC tumors showed stronger β-catenin activation upon transformation, indicating divergent oncogenic mechanisms. These findings suggest that the proFT may act as a primary precursor site for UTJC, rather than a secondary metastatic location. Further studies should prioritize investigating proFT-specific molecular drivers and risk factors underlying UTJC and EC development. Such insights could pave the way for precision prevention strategies, particularly in high-risk populations.

In conclusion, our findings highlight the critical role of regional cell lineages in tumor development and offer valuable insights for the development of precision therapies targeting these specific lineages.

## Methods

### Patient samples

Human normal fallopian tube tissue samples, along with ovarian cancer (OC), proFTE tumor and endometrial carcinoma (EC) tissue samples, were collected from patients who underwent cancer surgery in the Department of Gynecologic Oncology at the Third Affiliated Hospital of Guangzhou Medical University. In addition, ovarian cancer tissue microarrays (biomax, Cat#OV991) were employed for immunofluorescence staining. The approval of the Ethics Committee of Guangzhou Medical University (No. 2025087). Detailed patient and clinical information can be found in Table S1. Fresh tissues were collected and placed on ice, shipping to the laboratory within 2 hours. These human samples were used for primary organoids generation, immunostaining assays, or scRNA-seq analysis.

### Mice

All animal experiments were approved by the Guangzhou Medical University with project license of GY2022-218. Tp53^-/-^; Kras^G12D^; Cas9^l/l^ (PKC) mice were gifted from Xiaodong Wang lab (NIBS, Beijing) and maintained in a specific-pathogen-free facility (12hour (h) light / dark cycle, temperature 21–23 °C, humidity 30–70 %). PKC mice were used to construct distal and proximal fallopian tube derived organoid models. Wild type C57/BL6 mice and nude mice were purchased from Guangdong Medical Laboratory Animal Center, wild type C57/BL6 mice were used to construct distal and proximal fallopian tube derived organoid models. Female C57/BL6 mice aged 6 weeks with normal estrous cycles were included; mice with overt ovarian pathologies or systemic diseases were excluded. Nude mice and C57/BL6 mice were used to examine the difference in growth of PKP disFTO-driven and PKP proFTO-driven tumors via mammary fat pad injection or orthotopic injection. No animals were lost during the experiments; all mice survived until the scheduled endpoint. Only female mice were used in this study as specified in the protocol.

### Organoid cultures and assays

The *in vitro* organoids were generated with similar protocol as described before ^23, 49^. Briefly, the distal and proximal fallopian tubes of human and mice were separately harvested under a microscope and subsequently minced and digested with enzyme (Collagenase / Dispase®, Roche, cat#10269638001) at 37 °C for 1 h. The digested samples were centrifuged at 600xg for 5 min, and subjected to a second digestion by TryPLE (gibco, cat#12604013) at 37 °C for 10 minutes (min). DMEM (CORNING, cat#10-013-CVRC) with 1% FBS (Lifeman, cat#FBS001S) was added to terminate digestion, and cells were passed through a 70 μm strainer (Corning, cat#352350), centrifugation for 5 min at 600×g, washing one time with Ad+++ medium ((Advanced DMEM/F-12 (Thermo Fisher, cat#12634028), 100X HEPES (Coolaber, cat#SL6051-500), 100X GlutaMAX™ (gibco, cat#35050061), 100X Penicillin-Streptomycin (Gibco, cat#15140122)). Fallopian tube cells were seeded in 50μl marigel (Croning, cat#356231) mixed 1:1 with Ad+++ medium in 24-well plates (Croning, cat#3524). The organoids culture medium includes Ad+++, 50ng/mL Recombinant murine Wnt-3a (PeproTech, cat#315-20-10), 100ng/mL Human R-spondin-1 (PeproTech, cat#120-38-20), 50ng/mL Recombinant Human NOGGIN (Peprotech, cat#120-10c-250), 500nM A8301 (WAKO, cat#035-24113), 10μM Y27632 (selleckchem, cat#S1049), 1.25mM N-Acetyl-L-cysteine (Sigma, cat#A9165), 10nM Nicotinamide (Sigma, cat#N0636), 50ng/ml Recombinant human EGF (PeproTech, cat#AF-100-15). The number of cells in each passage was determined by Taipan (invitrogen, cat#T10282) blue staining to ensure the de reproducibility of organoid growth. The Tdiameter of organoids was counted by image J after 5-6 days culture (n=3).

### SgRNA design, Lenti-virus production and Infection

The mouse sgRNA was designed according to the method of web at https://portals.broadinstitute.org/gpp/public/. Among the results shown in the web, one sgRNA against Pten were selected as shown in Table S2. The SgRNA was respectively constructed into the lenti-Guide-Puro plasmid for lenti-virus production and cell infection. Lenti-virus production was performed as reported before ^50^. Briefly, a mixture of the constructed plasmids and viral coat plasmids (pPAX2 and PMD2.G) were mixed in a ratio of 4:3:1. The mixture were transfected into 293T cells with jetPRIME (polyplus, cat#114-15) reagent. The viral supernatant of 293T cells were harvested 72 h after transfection, subsequently passed through a 0.45 μm filter and concentrated with TransLvTM Lentivirus Precipitation Solution (transgen, cat#FV101-01). The 293T cell line (RRID:CVCL_0063) was authenticated by short tandem repeat profiling and tested negative for mycoplasma contamination.

For lentivirus infection of organoids, the oragnoids were digested into single cells, and the cells were resuspended with 500μL Ad+++ medium in an ultra-low adsorption 24-well plate (corning, 3473). 5μg/ml Polybrene (Beyotime, cat#C0351) was added to each well to help increase lentivirus infection efficiency. The 24-well plate was centrifuged at 600×g for 1 h at 37°C, and subsequently placed in an incubator at 37°C for 4-6h. After the lentivirus infection, the cells were resuscitated 50μl marigel mixed 1:1 with Ad+++ in 24-well plates. 3 days after lentivirus infection, 10 μg/ml puromycin (gibco, cat#A1113802) was added to the cultured organoids to screen for successfully infected organoids respectively. The knockout efficiency of organoids induced by sgRNA was detected by DNA extraction (Genomic DNA Small Volume Extraction Kit, Beyotime, cat#D0063) and PCR assay. The PCR primers were shown as Table S2.

### *In vitro* drug screen

PKP dFTO and PKP pFTO organoids were digested and seeded into 96-well plates at 1000 cells per well in gel free condition with 2D cell medium, which contained 1% Penicillin-Streptomycin, 4% FBS, 10ng/ml Recombinant human EGF (PeproTech, cat#AF-100-15) and 10ng/mL Recombinant Human Insulin (Pricella, cat#PB180432) in DMEM (CORNING, cat#10-013-CV). The 2D cell culture models were successfully converted in 96-well plates. After one day culture, the indicated concentrations of carboplatin, cisplatin, gemcitabine, niraparib, olaparib, ruaparib, doxorubicin, and paclitaxel were added in triplicate.

Drug susceptibility testing was performed as previously reported ^51^. Briefly, the drug dilution protocol was a fourfold gradient dilution method. After five days of culture, cell viability was measured with EnSight™ Multimode Microplate Reader (PerkinElmer) and PrestoBlue™ Cell Viability Assay Reagent (Thermo Fisher, cat#A13262) based on manufacturer instructions.

### Assessment of WNT growth factor dependency

To detect the dependence of proFTO and disFTO on WNT growth factor, organoids medium containing Wnt-3a, R-spodin1, noggin, EGF (WRNE-full) and organoids medium without Wnt-3a, R-spodin1, noggin, EGF (WRNE-minus) were prepared. According to the above passage method of organoids, 1×10^4^ cells each well were seeded in a 24-well plate. WRNE-full medium and WRNE-minus medium were respectively added to different wells. After 5 days of culture, the dependence of the proFTO and disFTO organoids on WNT growth factors were confirmed by detecting and counting the diameters of the organoids treated with different medium. The WNT growth factor-dependent experiment was repeated three times (n=3). Images were captured by using ZEISS microscope (Axiocam 208 color).

### ATAC-sequencing, data processing and bioinformatic analysis

ATAC-seq libraries were prepared using the Hyperactive ATAC-Seq Library Prep Kit for Illumina (Vazyme, cat#TD711-01) using the manufacturer’s instructions. About 100,000 cells were washed with precooled TW Buffer twice and lysed with precooled Lysis Buffer in PCR tube. Cells were cleaved to ice for 5 min and centrifuged at 500xg at 4℃ for 10 min. Cells were suspended in a 50μL reaction buffer containing 16.5μL TW Buffer, 0.5μL 10% Tween20, 0.5μL 1% Digitonin, 18.5μL Nuclease-free ddH2O, 10μL 5xTTBL, 4μL TTE Mix V50. The fragmentation reaction was carried out at 37°C for 30 min on PCR apparatus. Then, 5 μl Stop Buffer was added to the PCR tube and left for 5 min at room temperature to end the reaction. DNA was isolated immediately using ATAC DNA Extract Beads to generate 20μL fragmentation DNA. The resulting DNA was PCR amplified for 11 cycles with corresponding barcoding primers. The PCR amplified products were sorted using ATAC DNA Clean Beads in a 0.55× / 1.2× two-step process. After purification, the libraries were sequenced by Novogene company. For ATAC-seq data analysis, Fastp v0.20.0 was utilized to remove adapter sequences and low-quality reads. Paired-end reads were aligned using Bowtie2 v2.3.4.3 with the option --end-to-end --sensitive to ensure accurate and sensitive alignment. Duplicate reads were removed with Picard v2.18.17 using the parameter REMOVE_DUPLICATES=true. Peak calling was performed using SEACR v1.3 with a threshold of 0.01. Scatterplots, correlation plots, and visualization of peak files were generated using deepTools v2.27.1. Differential accessibility between conditions was evaluated with the DESeq2 R package, while peak annotation was conducted using the ChIPseeker v1.12.1 R package. To identify transcription factor binding sites, TOBIAS (Transcription factor Occupancy prediction By Investigation of ATAC-seq Signal) was employed ^52^. Specific peaks were defined as those with an M-value > 0.2 and a P-value < 0.05. Finally, ATAC-seq reads were visualized using IGV v2.18.4.

### CUT&Tag-sequencing, data processing and bioinformatic analysis

The CUT&Tag assay using the NovoNGS® CUT&Tag 4.0 High-Sensitivity Kit (Novoprotein, Cat#N259-YH01) was performed according to the manufacturer’s instructions. About 100,000 cells were subjected to formaldehyde cross-linking and resuspended in Dilution buffer. An appropriate amount of concanavalin A-coated magnetic beads (ConA beads) (10 μL per sample) were added to the binding buffer and washed once in the same buffer. Each time, the beads were placed on a magnetic rack to separate them from the buffer and then resuspended in binding buffer. The liquid was removed using a magnetic rack, and the cells were added to ConA beads. The mixture was incubated at a slow speed and room temperature for 10 min to allow the cells to bind to the beads. After cell-bead binding, the beads were collected using a magnetic rack, and the diluted primary antibody against H3K27ac (Abcam, cat#Ab4729) was added and incubated 2 h at room temperature.

The supernatant was removed, and the diluted secondary antibody was added and incubated at slow speed and room temperature for 1 h. The supernatant was removed, and the diluted transposase-containing buffer was added and incubated at slow speed and room temperature for 1 h to allow transposase-cell binding. The supernatant was removed, and the buffer containing MgCl_2_ was added. The mixture was incubated at slow speed and 37 °C for 1 h to fragmentize the DNA. To stop the fragmentation reaction, 5μL of Stop Buffer and 1 μL of Proteinase K were added to the incubated samples. The samples were vortexed at high speed and incubated at 55 °C in a metal bath for 10 min. Then, 2 times the volume of Tagment DNA Extract Beads (∼144 μL) was added to the terminated samples, mixed by pipetting, and incubated at room temperature for 5 min to extract the DNA fragments. The extracted DNA fragments were used as templates for PCR amplification and library construction. The i5 primer, i7 primer, template, and PCR enzyme from the kit were added, and the number of PCR cycles was determined based on the cell quantity and target protein selection. The PCR products were purified using NovoNGS® DNA Clean Beads. After purification, the libraries were sequenced by Novogene company. The analysis of cut&tag data was performed similar to ATAC-seq data as described before. We utilized IGV to visualize the CUT&Tag-seq reads and demonstrate the H3K27ac modification on target genes.

### Sample preparation and single-cell RNA sequencing

The procedure for digesting tissue into single cells was performed as previously described ^53^. Fresh tumor tissue was first mechanically minced into as small pieces as possible using a scalpel. Subsequently, the minced tissue was enzymatically digested in 10 ml of digestion medium consisting of 2 U/ml Dispase II (Sigma, D4693) and 2 mg/ml collagenase V (Sigma, C9263) in 10% FBS. The digestion mixture was gently shaken at 37 °C for 1 hour, with the contents pipetted up and down every 20 minutes to ensure thorough dissociation. After digestion, the cell suspension was filtered through a 40 μm cell strainer (Biosharp, cat#352340) to obtain a single-cell suspension. The single-cell suspension was centrifuged at 600xg for 5 minutes, and the supernatant was carefully discarded. The cell pellet was resuspended in erythrocyte lysis buffer (BOSTER, cat#AR1118) and incubated on ice for 5 minutes to lyse red blood cells. The lysis reaction was then quenched by adding D-PBS (Biosharp, cat#02-023-1ACS). Following another centrifugation at 600 xg for 5 minutes, the supernatant was removed, and the cell pellet was resuspended in D-PBS. Cell viability was assessed using trypan blue staining, with the percentage of viable single cells exceeding 80% of the total population. The resulting single-cell suspension was transported to AccuraMed company within 30 minutes and processed for single-cell isolation using the 10X Chromium Single Cell Instrument (10X Genomics). Subsequent steps, including barcoding, cDNA synthesis, and library preparation, were conducted according to the manufacturer’s standard protocols. High-quality libraries were selected for sequencing on the Illumina platform using PE150 configuration. scRNA-seq data analysis were performed as described previously ^40^. Briefly, single-cell RNA-seq reads were processed by the CellRanger (4.0.0) pipeline with default and recommended parameters. The Seurat (4.3.0) toolkit was employed for quality control. High-quality cells (genes per cell >500 and the content of mitochondrial genes <5%) were subjected to subsequent analyses. 2000 highly variable genes were selected, and the top 50 principal components were calculated using the PCA function.

The dimensionality reduction and cell clustering were performed using UMAP, and cell types were annotated according to well-known marker genes.

### EMT inhibitor-1 treatment

According to the above passage method of organoids, 1×10^4^ cells each well were seeded in a 24-well plate. The organoids and PKP proFTO cells were treated with 10 μM and 20 μM of EMT inhibitor-1 (MCE, cat#HY-101275), while 0.1% DMSO served as a control. After 5 days of treatment, the results were analyzed through images capturing, cell counting, RT-qPCR analysis and western blotting. The experiment was repeated three times (n=3).

### *In vivo* transplantation assays

For mammary fat pad injection and orthotopic injection, the manipulation procedure was performed as described before ^54^. PKP organoids were released, and 1×10^5^ cells were injected per nude mouse (n=2) by mammary fat pad injections. All the tumors in nude mice were removed half a month after injection. Moreover, PKP organoids were released, and 3×10^5^ cells were injected into one side of the ovary of each 6-week-old PKC homozygous mouse (n=3) by orthotopic injections. All the tumors in PKC homozygous mice were removed one month after injection.

### RNA-sequencing, data processing and bioinformatic analysis

Total RNA was extracted from organoids and tumors by RNAiso Plus (TaKaRa, cat#9109) according to manufacturer instructions, and RNA concentration and purity were determined using a NanoDrop 2000c (Thermo Scientific, Germany). RNA libraries were prepared and sequenced on Illumina NovaSeq 6000 sequencing platform using PE150 mode by BerryGnomics company.

Raw FASTQ of RNA-seq data sequencing files were aligned to mouse reference genome (build mm10) with bowtie2 (version 2.3.2) and tophat2 (version 2.1). Expression levels as raw “counts” were calculated from aligned reads with mapping quality ≥10 using htseq-count (version 0.6.0). Differential gene expression analyses were performed using DESeq2 (version 1.44.0) in R (R version 4.4). The differential expression genes with |log2FC|>1 and pvalue<0.05 were selected for GO analysis, KEGG analysis, and Venn plot analysis.

### RT-qPCR Analysis

The RNA extracted from organoids were reversed with HiScript III RT SuperMix for qPCR (+ gDNA wiper) (Vazyme, cat#R323). Quantitative reverse transcription polymerase chain reaction (RT-qPCR) was then performed on a QuantStudio™ 1 Plus instrument using specific primers and ChamQ Universal SYBR qPCR Master Mix (Vazyme, cat#Q711-02). RT-qPCR results were calculated by delta-delta-Ct values. The RT-qPCR primers used in this study are shown as Table S3.

### Western blot

The organoids were released from matrigel with cold PBS, organoids were cleaved with RIPA lysate (Beyotime, cat#P0013K) containing 100× protease inhibitors (Sigma, cat#p8849) and 100× phosphatase inhibitors (merck, cat#524625-1SET). Protein concentrations were measured using protein standards (Thermo, cat#23209) and BCA Protein Concentration Assay kit (Beyotime, cat#P0012). The proteins were separated by 10% sodium dodecyl sulfate-polyacrylamide gel electrophoresis, and then transferred to 0.22μm PVDF membranes (Millipore, cat#ISEQ00010) that then was blocked in 5% skim milk (APPLYGEN, cat#P1611-500) / TBS (Roles-Bio, cat#RBG6-1) containing 0.1% Tween-20 (Solarbio, cat#T8220) for 1 h. The primary antibodies were added overnight at 4℃ and then washed with TBST, incubated with HRP-labeled secondary antibodies for 1h at room temperature, followed by washing with TBST (Roles-Bio, RBG8-1). The WB results were detected by ECL chemiluminescence kit (Biosharp, cat#BL520A). Image was acquired using the fully automatic chemiluminescence imaging system system from SINSAGE’s ChampChemi^®^610. The primary antibodies include ERK2 (Santa Cruz Biotechnology, cat#sc1647, RRID:AB_627547), PTEN (Cell signaling technology, cat#9559S), Vim (Cell signaling technology, cat#5741S), and E-cad (Abcam, cat#Ab231303, RRID:AB_2923285). Secondary antibodies include HRP Goat Anti-Mouse IgG (H+L) Antibody (Apexbio, cat#K1221), HRP Goat Anti-Rabbit IgG (H+L) Antibody (Apexbio, cat#K1223). All uncropped and unprocessed scans of the western blots were supplied in Supplementary Fig. 7 in the Supplementary Information.

### Histology and immunostaining

Tissues were immersed in 4% PFA for 24 h, organoids were released from Matrigel and fixed in 4% PFA for 15 min and transferred to Histogel before tissue processing and embedding. Formalin-fixed, paraffin-embedded tissue sections (5μm) were stained with hematoxylin-eosin (H&E) or immunohistochemical staining. The slices were baked at 65°C for 1 h, de-paraffinized with xylene, rehydrated with gradient alcohol. For antigen retrieval, slides were autoclaved in 0.01 M citrate buffer (pH 6.0) for 15 min, and the endogenous peroxidase activity of the slices was quenched at 3% H_2_O_2_ for 10 min. The slices were washed with PBS for 10 min, blocked with 5% BSA (Solarbio, cat#A8020) for 1h at room temperature. Primary antibodies were added into the slices overnight at 4°C. Then the slices were washed in PBS for 10 min. Secondary antibodies with HRP-labeled were incubated at room temperature for 1h, and washed again with PBS. The signal was visualized using DAB color development kit (ZSGB-BIO, cat#ZLI-9019) according to the reagent manufacturer’s instructions, stained with hematoxylin for 3min, and peracid ethanol. Then the slices were treated by lithium carbonate to reverse blue for 30s, and then dehydrated with gradient alcohol, permeabilized with xylene, and sealed with neutral resin. Primary antibodies include: Wt1 (Abcam, cat#ab89901, RRID:AB_2043201), Pax8 (Proteintech, cat#10336-1-AP, RRID:AB_2236705), Acetylated-α-tubulin (Sigma, cat#T7451, RRID:AB_609894), CK7 (BOSTER Biological Technology, cat#BM1618), Ki67 (Arigo, cat#ARG11083), Stathmin1 (Proteintech, cat#11157-1-AP, RRID:AB_2197114). Secondary antibodies included HRP Goat Anti-Rabbit IgG (HRP Goat Anti-Rabbit, cat#K1223), HRP Goat Anti-Mouse (Apexbio, cat#K1221). Images were captured by using OLYMPUS microscope (BX43).

For immunofluorescence staining, organoids released from matrigel were fixed in 4% PFA for 15 min at room temperature and washed once with PBS. Membranes were permeabilized with PBS containing 0.5% TritonX-100 (Solarbio, cat#T8200) and 1% BSA (Solarbio, cat#A8020) for 30 min at room temperature. Then organoids were blocked with PBS containing 1% BSA and 0.1% Tween-20 for 1 h at room temperature. Primary antibody was added overnight at 4°C. The organoids were washed with PBS, and incubated with fluorescein-labeled secondary antibody for 1 h at room temperature. The plates were then washed with PBS, incubated with 0.1μg/ml DAPI (Sigma, cat#D9542) at room temperature for 10 min, and sealed by the addition of the anti-fluorescence quender Fluoromount-G (Southbiotech, cat#0100-01). Primary antibodies included Wt1 (Abcam cat#ab89901), Pax8 (Proteintech, cat#10336-1-AP), Acetylated-α-tubulin (Sigma, cat#T7451), Foxj1(Invitrogen, cat#14-9965-82), CK7 (Arigo, cat#ARG11083), Ki67 (Arigo, cat#ARG11083), Gpx3 (R&D Systems, cat#AF4199), β-Catenin (BD Transduction Laboratories™, cat#610154). Secondary antibodies included Alexa Fluor^®^ 488-AffiniPure Goat Anti-Mouse (jackson, cat#115-545-003), Alexa Fluor^®^ 488-AffiniPure Goat Anti-Rabbit (jackson, cat#111-545-003), Alexa Fluor^®^ 594-AffiniPure Goat Anti-Mouse (jackson, cat#115-585-003), Alexa Fluor^®^ 594-AffiniPure Goat Anti-Rabbit (jackson, cat#111-585-003), Cy5 Goat Anti-Mouse (ApexBio, cat#K1210), Cy5 Rabbit Anti-Goat (Apexbio, cat#K1216), Alexa Fluor® 647 Conjugate Anti-rat IgG (H+L) (Cell signaling technology, cat#4418S). Images were obtained by using Nikon confocal microscope.

### Quantification and statistical analysis

Image analysis and quantification was done using Image J software. Statistical analyses were performed, and graphs were generated using GraphPad Prism (version 9.5.1). To calculate the percentage of Vim^+^E-cad^+^ double-positive cells, we counted both the E-cad+Vim+ cells and the total number of E-cad-positive cells in at least five randomly selected fields (40x magnification) per sample. From this, we determined the percentage of double-positive cells among all E-cad-positive cells. For the β-catenin nuclear cells, we employed the same approach, examining at least five 40x fields per sample. We counted the β-catenin nuclear cells alongside the total number of β-catenin positive cells to compute the percentage of nuclear cells. Data are presented as mean ± SEM. P value less than 0.05 was considered statistically significant. Unless otherwise specified, all studies for which data are presented are representative of at least two independent experiments.

## Acknowledgements

We thank Dr. Lu Lv and Dr. Ze Zhang from Prof. Xiaodong Wang’s Lab (NIBS, Beijing) for kindly providing PKC mice. We also extend our gratitude to Prof. Dapeng Hao and Dr. Tianhao Li from Harbin University of Science and Technology for their valuable discussions and assistance with data analysis. LW and HW contributed equally to this work. Work on this project was supported by grants from the National Natural Science Foundation of China (82173370 to SZ), Science and Technology Program of Guangzhou (202201020573 to SZ), Victoria’s Secret Global Fund for Women’s Cancers Career Development Award, in partnership with Pelotonia & AACR (22-20-73-ZHAN to SZ).

## Author contributions

Conceptualization: SZ, ZT, WS. Methodology: LW, LH, JL, WL. Investigation: LW, ZH, YW, HW. Visualization: LW, LH. Funding acquisition: SZ. Supervision: ZT, SZ. Clinical analysis: SC, HW. Writing original draft: LW, SZ. Writing, review & editing: all authors.

---

**Supplementary Fig. 1.**
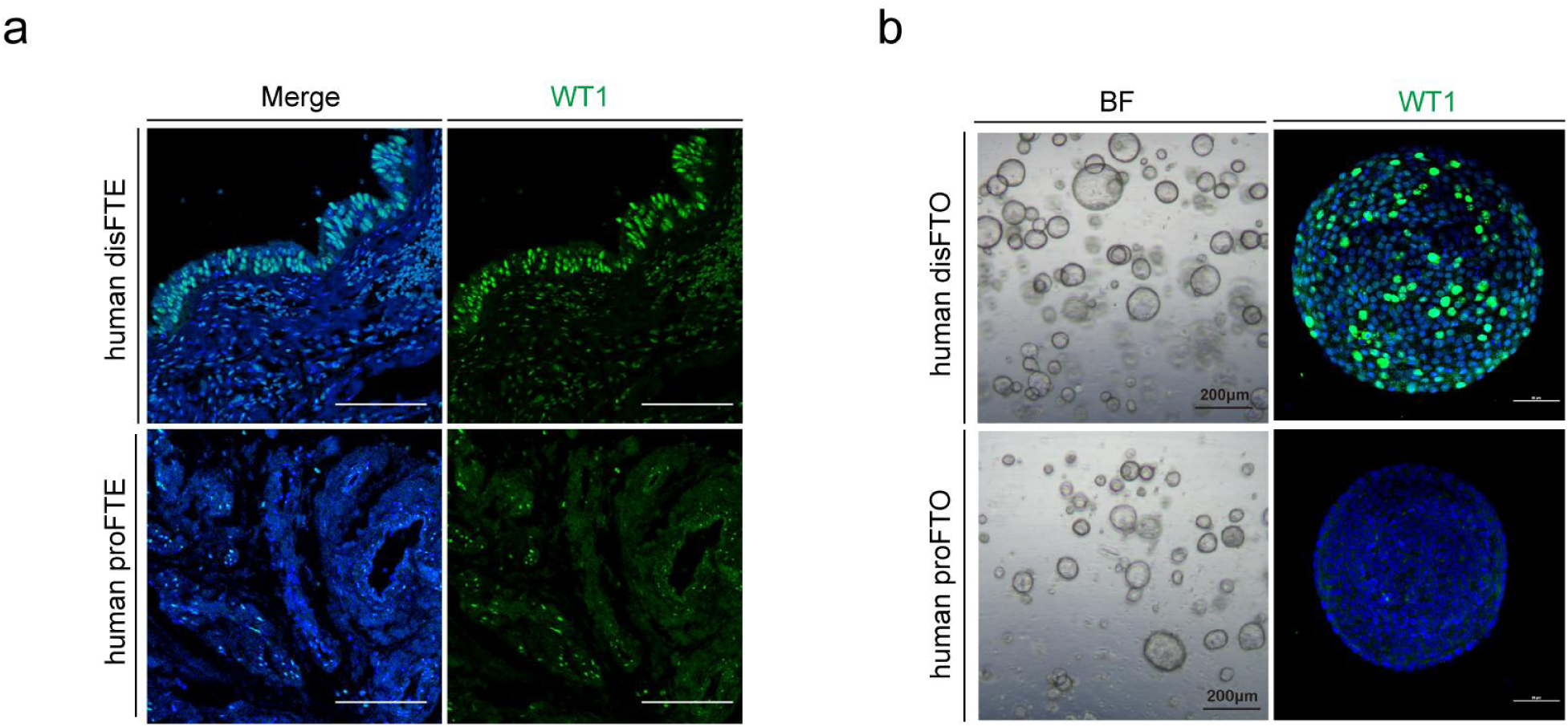
, related to Fig.1 | WT1 Expression Is Higher in Human disFTE Than proFTE and Recapitulated in Corresponding Organoids. a Representative Immunofluorescent staining of WT1 and DAPI in human disFTE and proFTE. Scale bars, 100 μm. b Representative Bright-field image and Immunofluorescent staining of WT1 and DAPI in human organoids cultured for 12 days. Scale bars:200 μm (left panel) and 50 μm (right panel).

**Supplementary Fig. 2.**
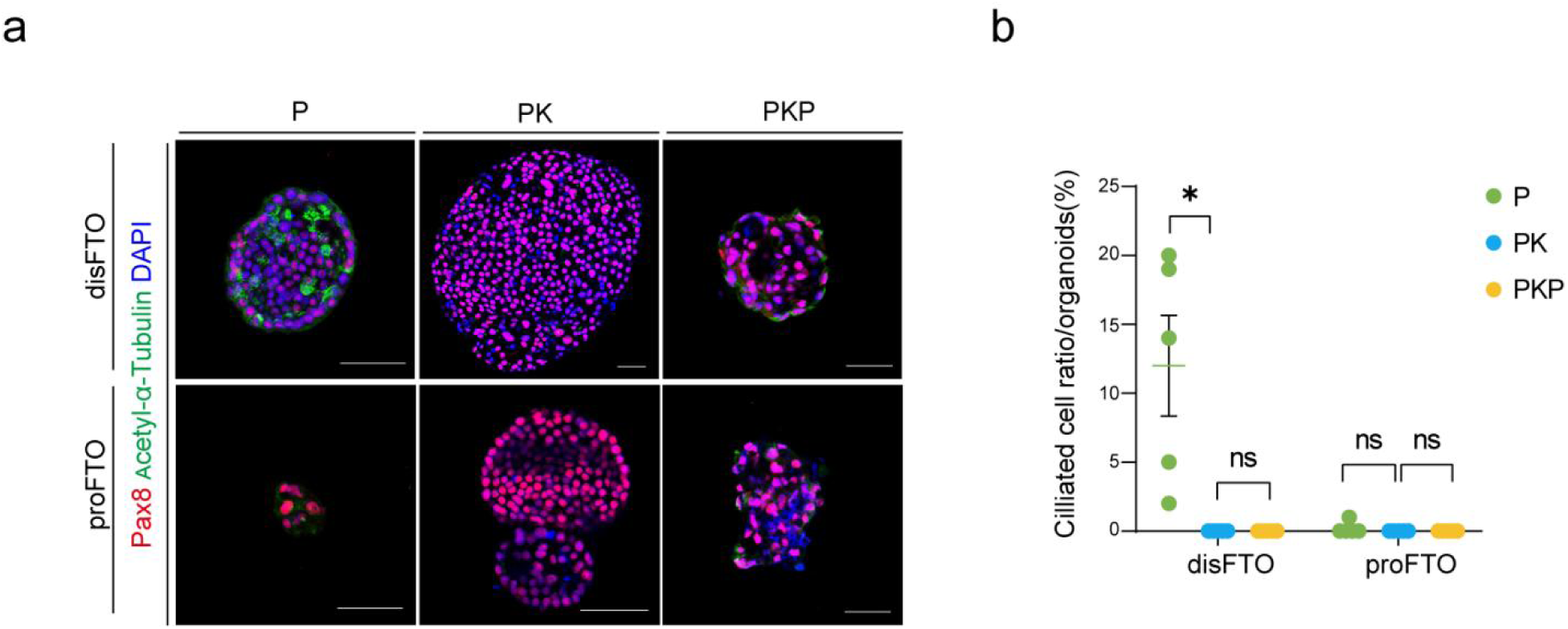
, related to. Fig. 2 **| Oncogenic Incorporation Blocks Ciliated Cell Differentiation in disFTO. a** Immunofluorescent staining of organoids showing the presence of Pax8, Acetyl-α-Tubulin, and DAPI. Scale bar: 50 μm. **b** Percentage of ciliated cells (Acetylated Tubulin+) in the indicated organoids. Organoids were calculated after 10 days in culture. Data represent mean ± SEM, *P < 0.01.

**Supplementary Fig. 3.**
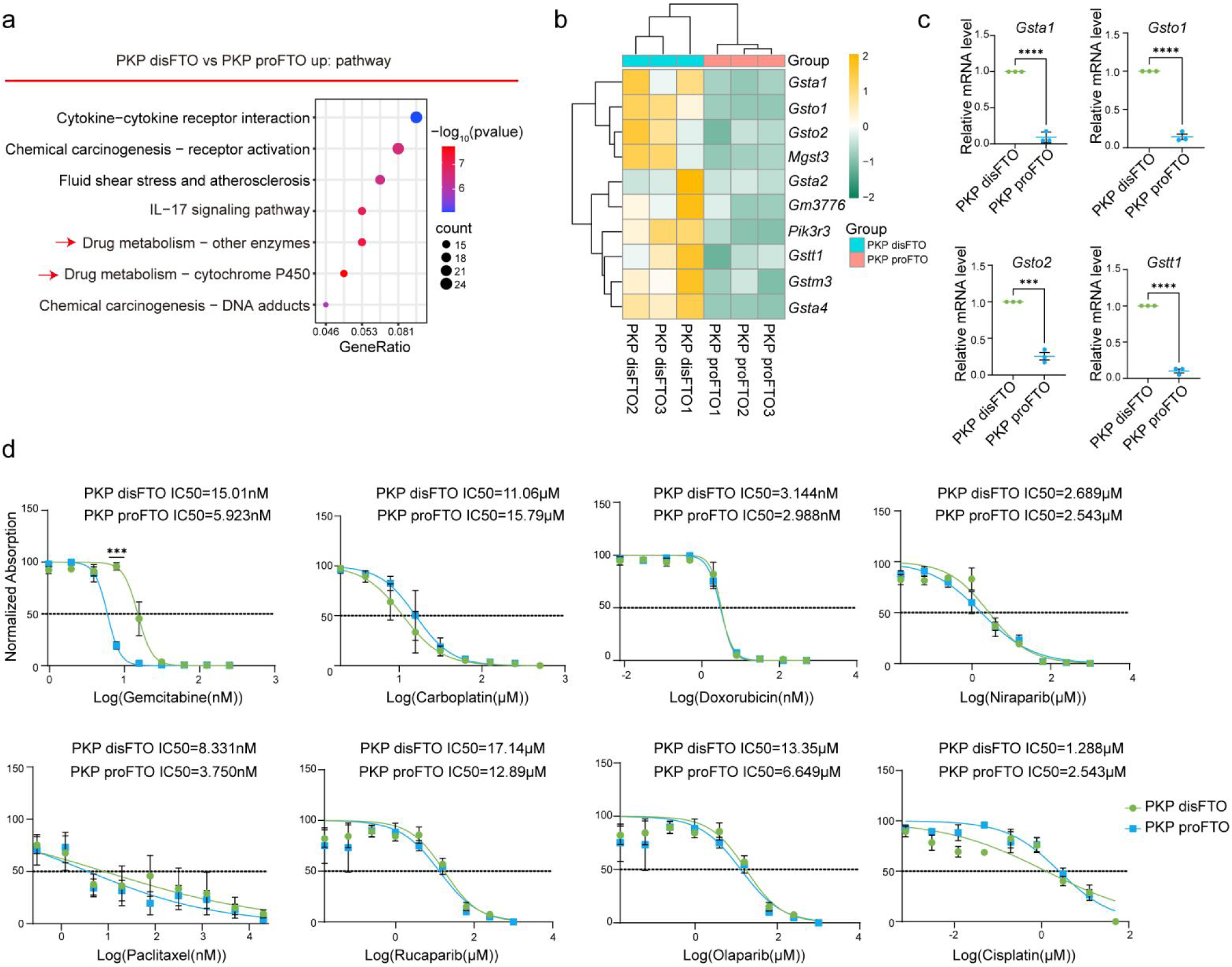
, related to. Fig. 3 **| Drug sensitivity analysis in PKP dFTO and pFTO organoids. a** Pathway analysis of upregulated genes in PKP dFTO compared to PKP pFTO. **b** Heatmap showing the expression of genes in pathways related to drug metabolism. **c** qPCR analysis of selected genes from the drug metabolism-related pathway. The scatter plot represents fold-changes relative to PKP dFTO, set as 1 (RQ, relative quantification). Error bars represent mean ± SEM. β-actin was used as an endogenous control. n = 3 independent experiments, with triplicates for each gene per experiment. NS: not significant; *P < 0.05; **P < 0.01; ***P < 0.001; ****P < 0.0001. **d** Drug dose-response curves for PKP dFTO and PKP pFTO treated with the following drugs: Gemcitabine (0–250 nM), Carboplatin (0–500 μM), Doxorubicin (0–500 μM), Niraparib (0–1 mM), Paclitaxel (0–20 μM), Rucaparib (0–250 μM), Olaparib (0–1 mM), and Cisplatin (0–1 mM). Cell viability was calculated relative to 0.01% DMSO-treated control cells. Data represents mean ± SEM. NS: not significant; *P < 0.05; **P < 0.01; ***P < 0.001; ****P < 0.0001. Source data are provided as a Source Data file.

**Supplementary Fig. 4.**
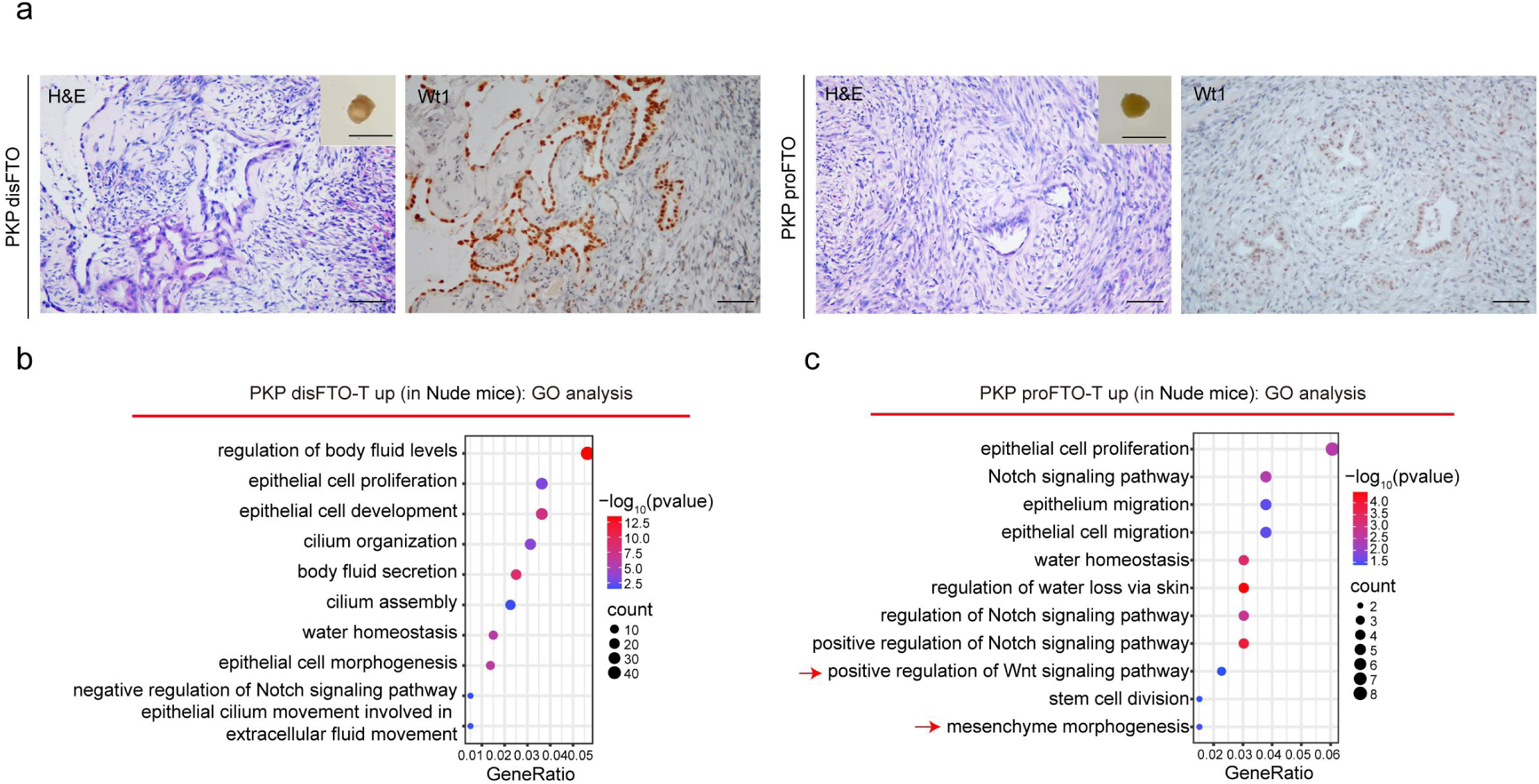
, related to. Fig. 4 **| proFTO and disFTO derived tumors Exhibit Distinct Morphology and WT1 Expression Patterns. a** Representative images of another organoid-derived tumor formed after mammary fat pad injection of PKP disFTO and PKP proFTO. Scale bars represent 1 cm. H&E and IHC staining of tumors formed by PKP dFTO and PKP pFTO, showing expression of WT1. Scale bars represent 50 μm. **b** GO enrichment analysis of upregulated genes in PKP disFTO-derived tumors (PKP disFTO-T) compared with PKP proFTO-derived tumors (PKP proFTO-T) in nude mice. **c** GO enrichment analysis of upregulated genes in PKP proFTO-T compared with PKP disFTO-T in nude mice.

**Supplementary Fig. 5.**
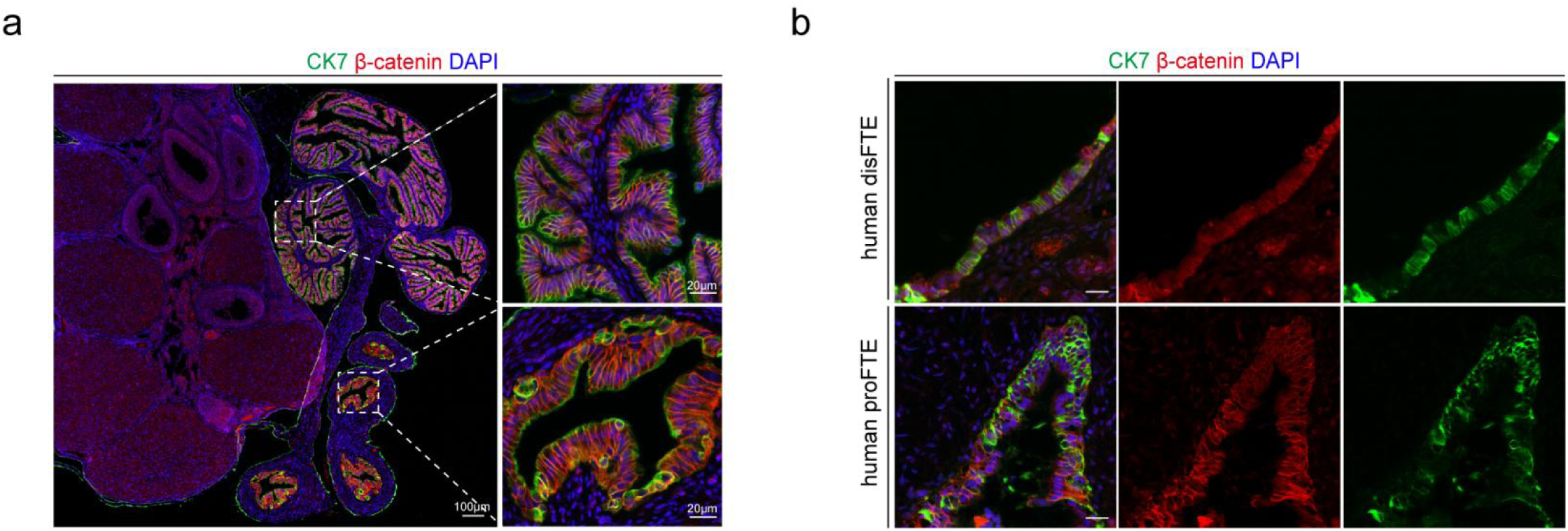
, related to. Fig. 4**, 7 | Compartment-Specific β-Catenin Expression in Normal Fallopian Tube Epithelium. a** Immunofluorescence staining of CK7, β-catenin, and DAPI in C57BL/6 mouse fallopian tube epithelium (FTE). Scale bars: 20 μm and 100 μm. **b** Immunofluorescence staining of human disFTE and proFTE with CK7, β-catenin, and DAPI. Scale bar: 20 μm.

**Supplementary Fig. 6.**
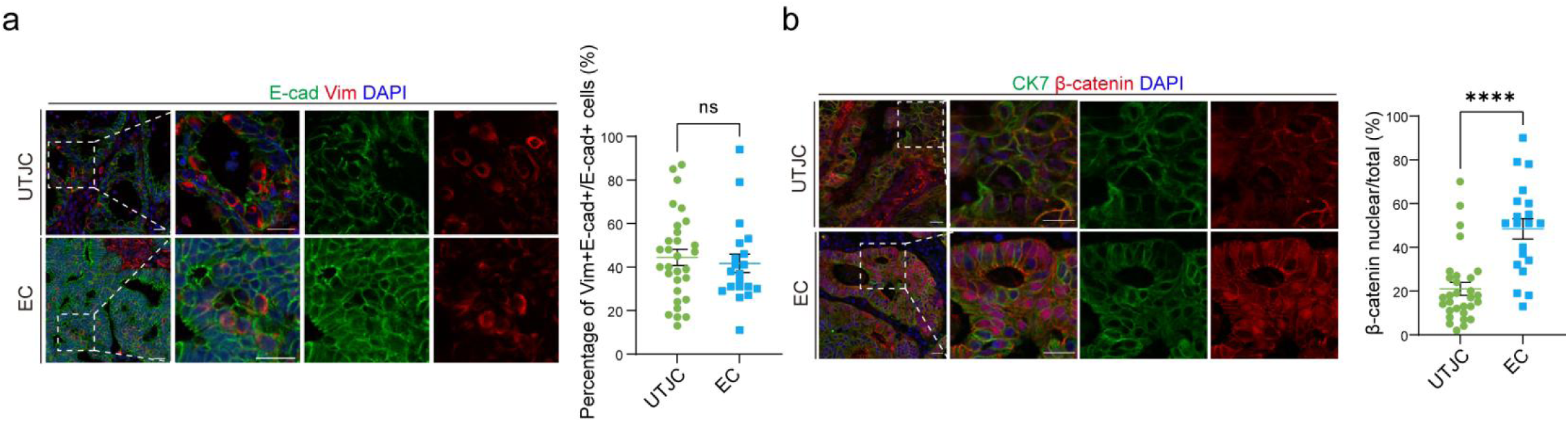
, related to. Fig. 7 **| The Proportion of E-CAD⁺Vim⁺ Double-Positive Cells and β-Catenin Nuclear-Localized Cells in EC. a** Immunofluorescence staining of UTJC, and EC with E-cad, Vim, and DAPI (n = 4 – 6 patient samples, count 5 fields of vision for each patient). Scale bar: 20 μm. The percentage of E-cad and Vim double-positive cells among E-cad-positive cells in human UTJC, and EC is presented as mean ± SEM (unpaired t-test, ns: not significant). **b** Immunofluorescence staining of UTJC, and EC with CK7, β-catenin, and DAPI (n = 4 – 6 patient samples, count 5 fields of vision for each patient). Scale bar: 20 μm. The percentage of nuclear β-catenin-positive cells among total β-catenin-positive cells in human UTJC, and EC is presented as mean ± SEM (unpaired t-test, *P < 0.05).

**Supplementary Fig. 7.**
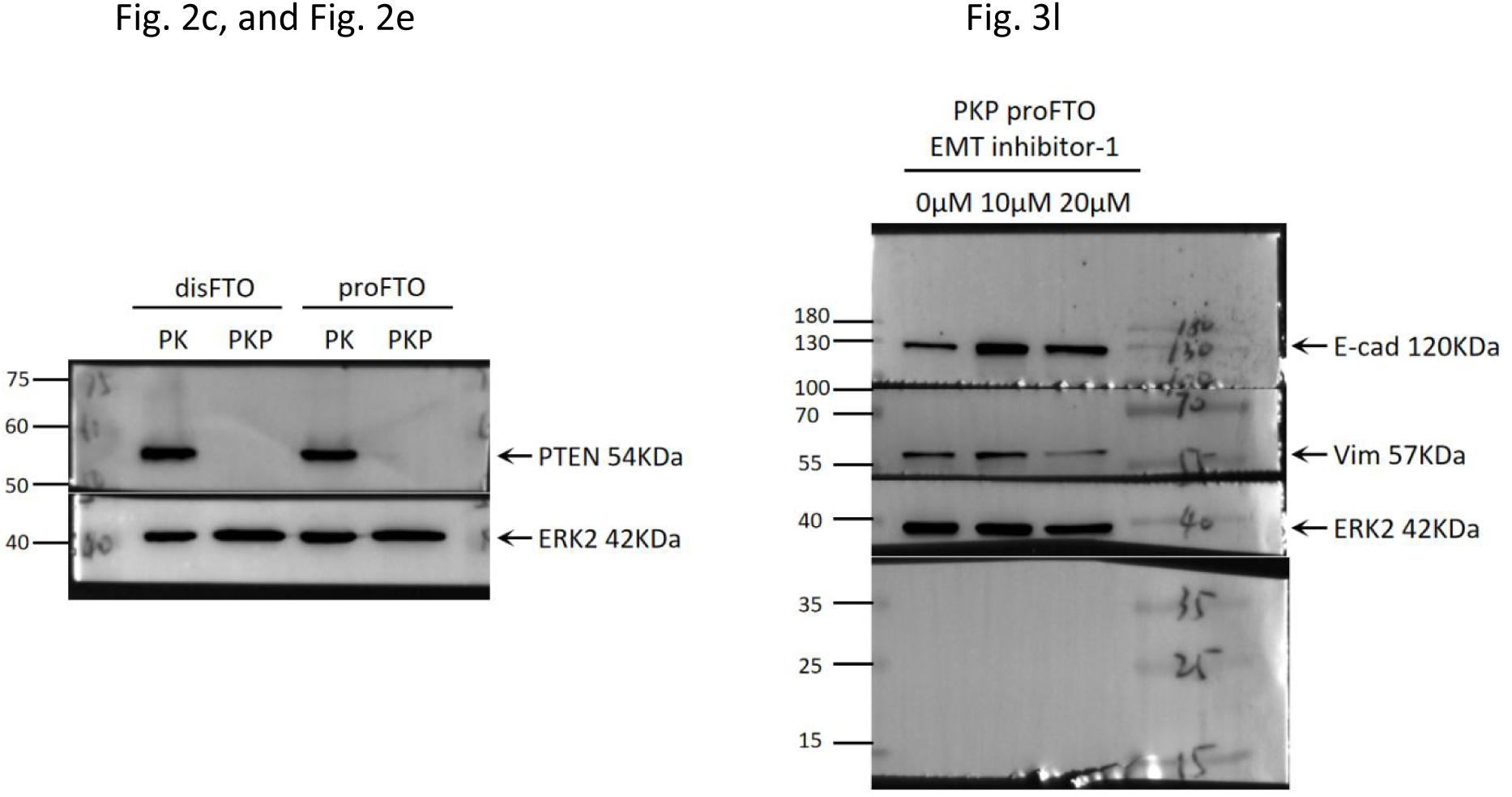
| Unprocessed images of immunoblotting.

**Table S1.** Clinical Information of Patient Samples Used in the Study.

| Patient ID | Age | Category | Other information | Purpose |
| --- | --- | --- | --- | --- |
| OV991 C7 | 63 | HGSC | High grade serous carcinoma stage I | IF |
| OV991 C8 | 51 | HGSC | High grade serous carcinoma stage I | IF |
| OV991 C10 | 47 | HGSC | High grade serous carcinoma stage IA | IF |
| OV991 D2 | 54 | HGSC | High grade serous carcinoma stage IA | IF |
| OV991 D3 | 56 | HGSC | High grade serous carcinoma stage IA | IF |
| OV991 D4 | 70 | HGSC | High grade serous carcinoma stage IB | IF |
| OV991 D5 | 36 | HGSC | High grade serous carcinoma stage II | IF |
| OV991 D6 | 66 | HGSC | High grade serous carcinoma stage II | IF |
| OV991 D7 | 62 | HGSC | High grade serous carcinoma stage IIB | IF |
| OV991 D8 | 50 | HGSC | High grade serous carcinoma stage IIIA | IF |
| OV991 D9 | 75 | HGSC | High grade serous carcinoma stage III | IF |
| OV991 E1 | 49 | HGSC | High grade serous carcinoma stage IIIA | IF |
| OV991 E3 | 48 | HGSC | High grade serous carcinoma stage IIIB | IF |
| OV991 E4 | 64 | HGSC | High grade serous carcinoma stage IIIC | IF |
| OV991 E5 | 52 | HGSC | High grade serous carcinoma stage IV | IF |
| 393717 | 36 | EC | Endometrioid adenocarcinoma | IF |
| 396100 | 41 | EC | Endometrioid adenocarcinoma stage II | IF |
| 400493 | 58 | EC | Endometrial adenocarcinoma stage IA | IF |
| 2190086 | 73 | EC | Low-grade endometrioid carcinoma G1 | IF |
| 324903 | 53 | UTJC | Highly differentiated endometrioid adenocarcinoma stage IA,<br>cauliflower mass on the left uterine horn | IF |
| 355092 | 55 | UTJC | Endometrioid adenocarcinoma stage IIIC1,<br>there's a mass on the left uterine horn | IF |
| 441657 | 57 | UTJC | Endometrioid adenocarcinoma stage IB,<br>there's a mass on the left uterine horn | IF |
| 441586 | 75 | UTJC | Endometrioid adenocarcinoma stage IB,<br>cauliflower mass near the left uterine horn | IF |
| 373827 | 56 | UTJC | High-grade endometrioid adenocarcinoma stage IIIA,<br>there is a cauliflower mass in the inner membrane of the left uterine<br>horn | IF |
| 159703 | 63 | UTJC | Endometrial carcinoma stage IIC,<br>cauliflower mass was found near the right uterine horn | IF |
| 471235 | 39 | EC | Endometrioid adenocarcinoma stage IA | ScRNA-seq |

**Table S2.**
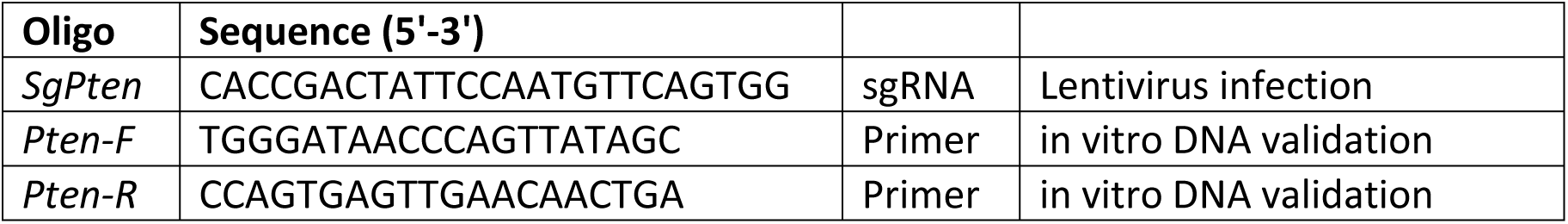
The List of Primers and DNA Validation Sequences Used for the Construction of SgRNA.

**Table S3.**
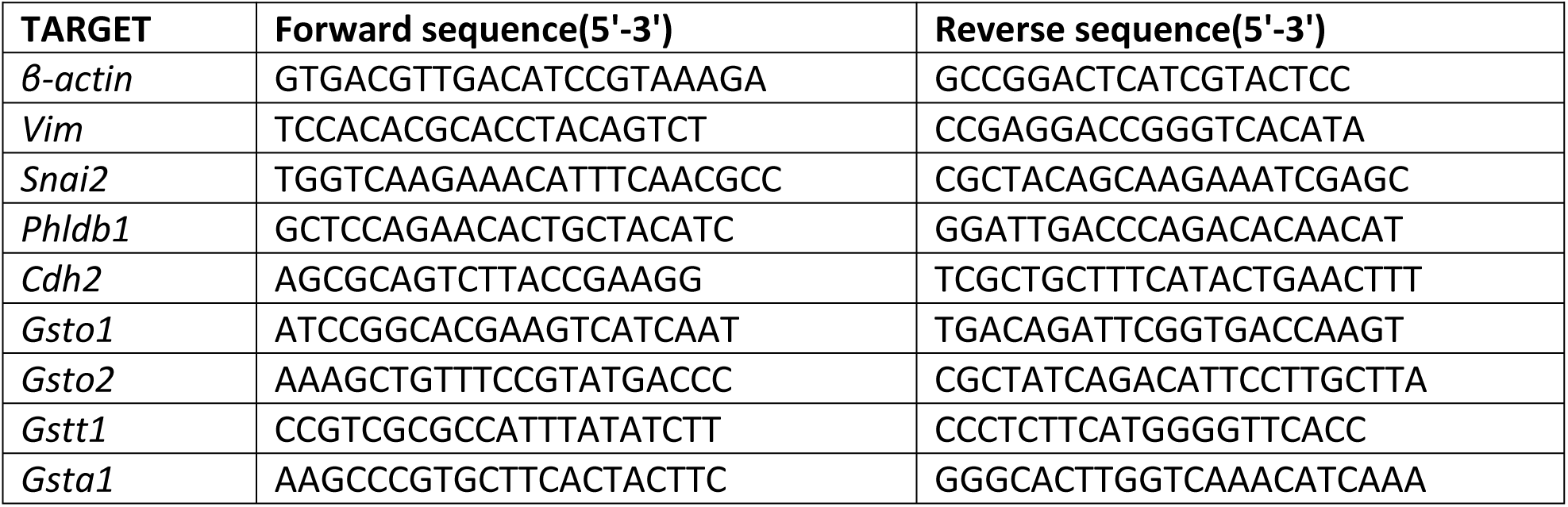
Primer List of RT-qPCR.

**Table S4.**
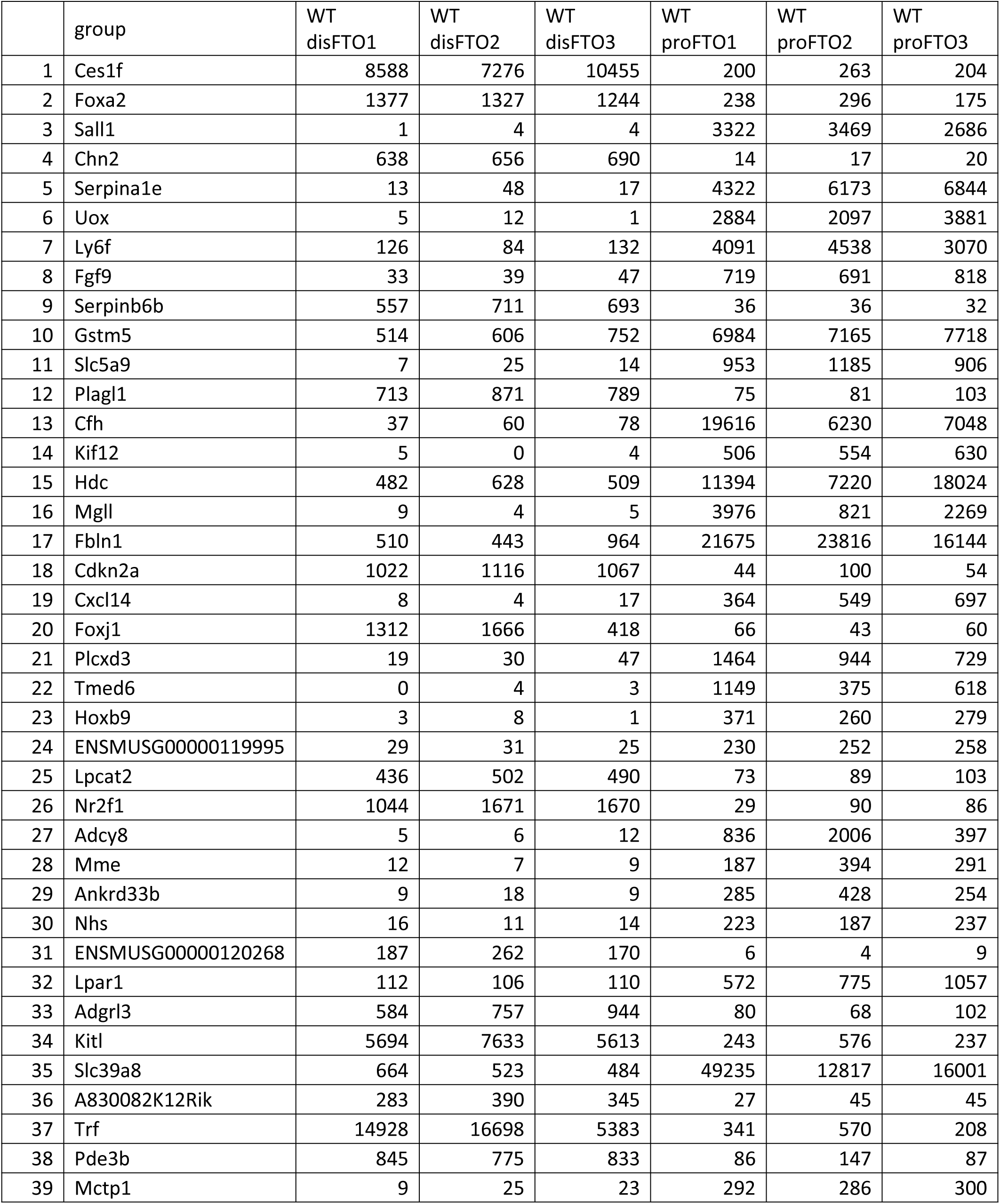
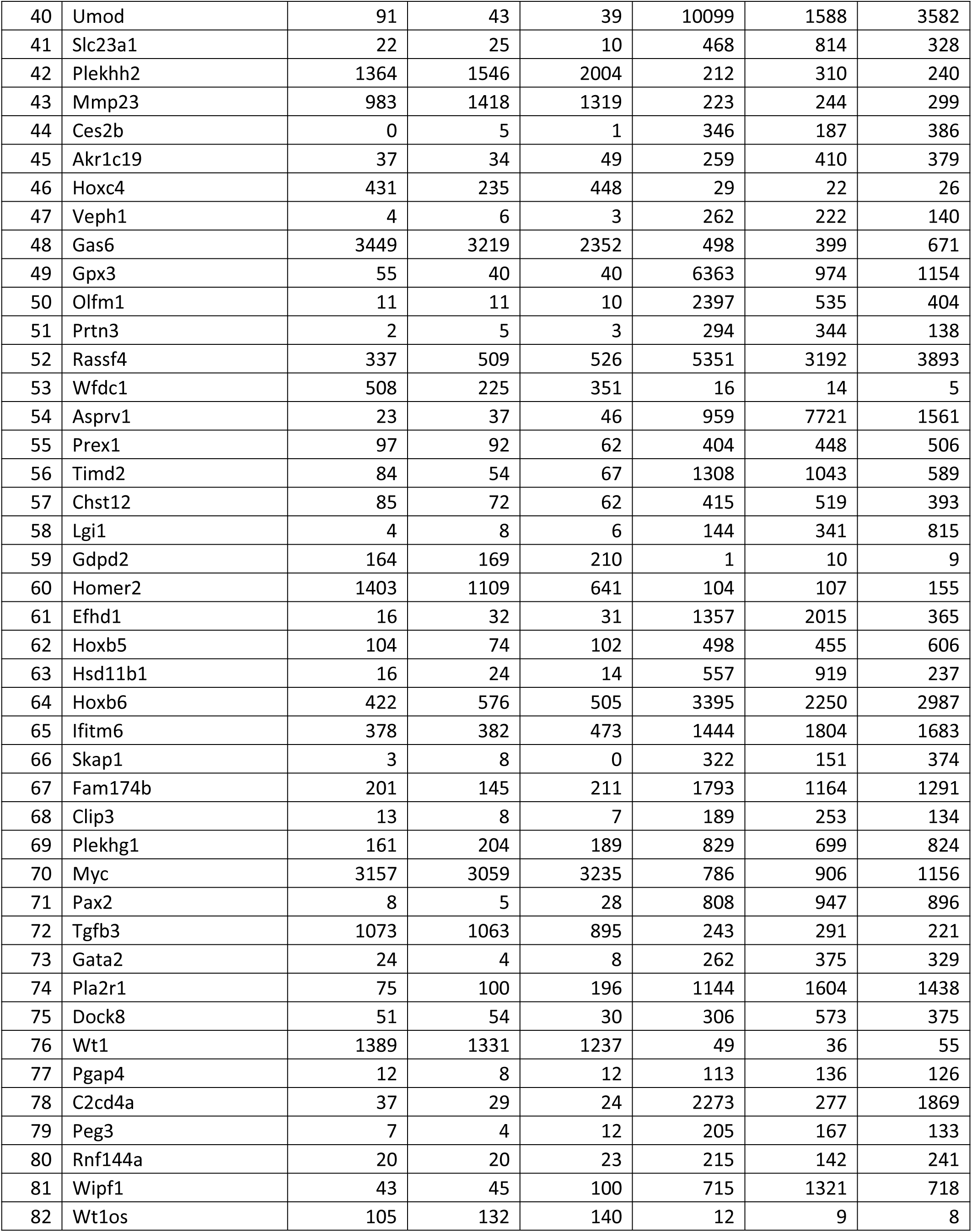
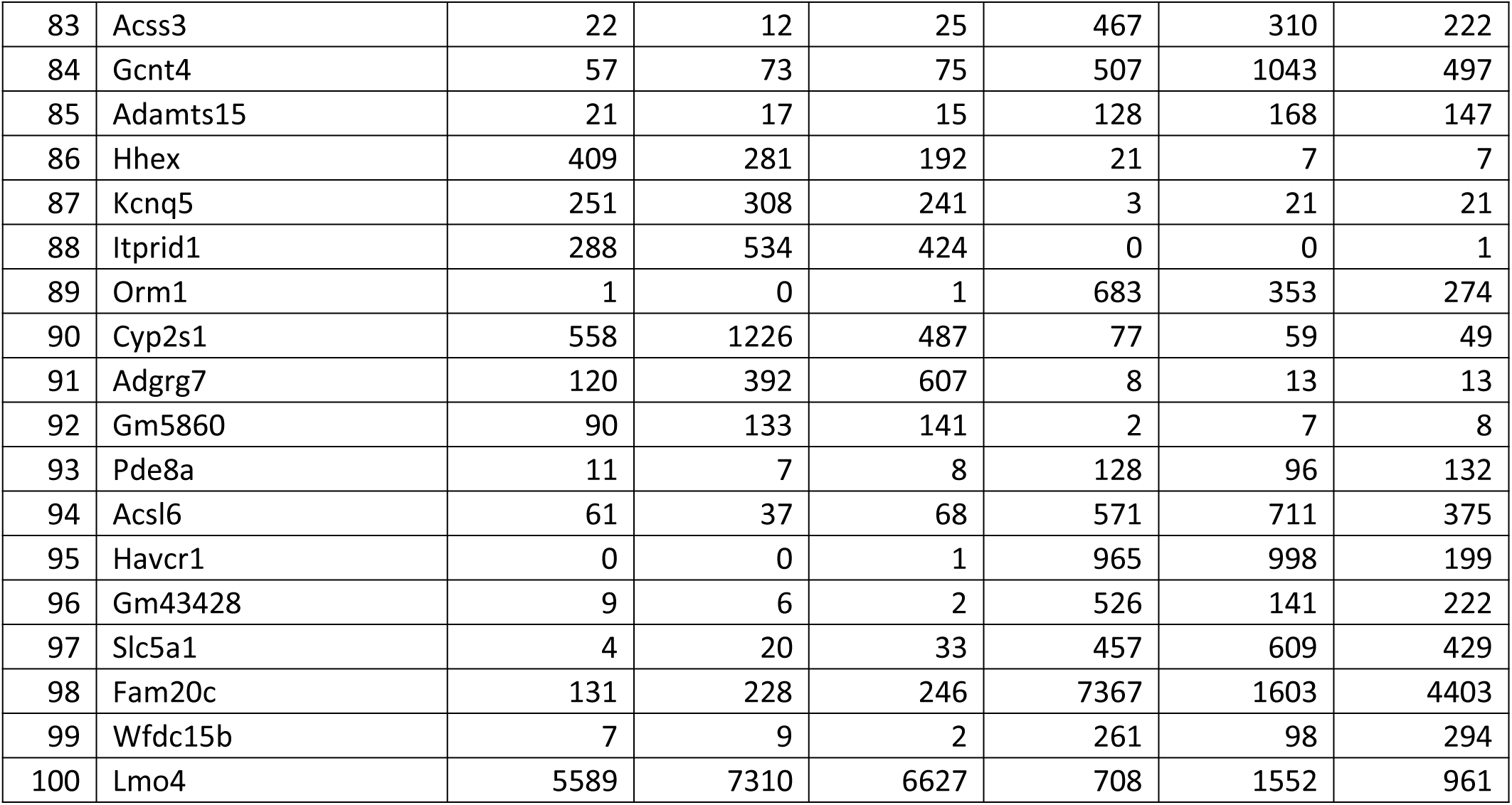
The Counts of Top 100 Differentially Expressed Genes Between WT disFTO and proFTO.

**Table S5.** Clinical Information of Patient Samples Used in the Study

| EMT_score gene | WNT_score gene |
| --- | --- |
| *VIM* | *FZD4* |
| *FN1* | *FZD8* |
| *OGN* | *PLPP3* |
| *TIMP3* | *TCF4* |
| *SPARC* | *TCF12* |
|  | *TNFRSF10* |
|  | *PRKAA1* |
|  | *LGR5* |
|  | *TMEM88* |
|  | *APCDD1* |

## References

1. Qin, G. et al. Distinct niche structures and intrinsic programs of fallopian tube and ovarian surface epithelial cells. iScience 26, 105861 (2023).

2. Yang, L. et al. Tissue-location-specific transcription programs drive tumor dependencies in colon cancer. Nat Commun 15, 1384 (2024).

3. Tatangelo, F. et al. Posterior HOX genes and HOTAIR expression in the proximal and distal colon cancer pathogenesis. J Transl Med 16, 350 (2018).

4. Ali, A. et al. Prostate zones and cancer: lost in transition? Nat Rev Urol 19, 101–115 (2022).

5. Missiaglia, E. et al. Distal and proximal colon cancers differ in terms of molecular, pathological, and clinical features. Ann Oncol 25, 1995–2001 (2014).

6. Eddy, C.A. & Pauerstein, C.J. Anatomy and physiology of the fallopian tube. Clin Obstet Gynecol 23, 1177–1193 (1980).

7. Tone, A.A. et al. The role of the fallopian tube in ovarian cancer. Clin Adv Hematol Oncol 10, 296–306 (2012).

8. Bast, R.C., Jr., Hennessy, B. & Mills, G.B. The biology of ovarian cancer: new opportunities for translation. Nat Rev Cancer 9, 415–428 (2009).

9. Roy, R., Chun, J. & Powell, S.N. BRCA1 and BRCA2: different roles in a common pathway of genome protection. Nat Rev Cancer 12, 68–78 (2011).

10. Hartmann, L.C. & Lindor, N.M. The Role of Risk-Reducing Surgery in Hereditary Breast and Ovarian Cancer. N Engl J Med 374, 454–468 (2016).

11. King, M.C., Marks, J.H., Mandell, J.B. & New York Breast Cancer Study, G. Breast and ovarian cancer risks due to inherited mutations in BRCA1 and BRCA2. Science 302, 643–646 (2003).

12. Eckert, M.A. et al. Genomics of Ovarian Cancer Progression Reveals Diverse Metastatic Trajectories Including Intraepithelial Metastasis to the Fallopian Tube. Cancer Discov 6, 1342–1351 (2016).

13. Lee, Y. et al. A candidate precursor to serous carcinoma that originates in the distal fallopian tube. J Pathol 211, 26–35 (2007).

14. Kuhn, E. et al. TP53 mutations in serous tubal intraepithelial carcinoma and concurrent pelvic high-grade serous carcinoma--evidence supporting the clonal relationship of the two lesions. J Pathol 226, 421–426 (2012).

15. Paik, D.Y. et al. Stem-like epithelial cells are concentrated in the distal end of the fallopian tube: a site for injury and serous cancer initiation. Stem Cells 30, 2487–2497 (2012).

16. Sowamber, R. et al. Integrative Transcriptome Analyses of the Human Fallopian Tube: Fimbria and Ampulla-Site of Origin of Serous Carcinoma of the Ovary. Cancers (Basel) 12 (2020).

17. Rose, I.M. et al. WNT and inflammatory signaling distinguish human Fallopian tube epithelial cell populations. Sci Rep 10, 9837 (2020).

18. Badiner, N., Nchako, C.M., Ma, L. & Frey, M.K. Ovarian cancer arising from the proximal fallopian tube in a patient with a BRCA2 mutation. Gynecol Oncol Rep 37, 100795 (2021).

19. MacLaughlin, D.T., Teixeira, J. & Donahoe, P.K. Perspective: reproductive tract development--new discoveries and future directions. Endocrinology 142, 2167–2172 (2001).

20. Ulrich, N.D. et al. Cellular heterogeneity of human fallopian tubes in normal and hydrosalpinx disease states identified using scRNA-seq. Dev Cell 57, 914–929 e917 (2022).

21. Ghosh, A., Syed, S.M. & Tanwar, P.S. In vivo genetic cell lineage tracing reveals that oviductal secretory cells self-renew and give rise to ciliated cells. Development 144, 3031–3041 (2017).

22. Dinh, H.Q. et al. Single-cell transcriptomics identifies gene expression networks driving differentiation and tumorigenesis in the human fallopian tube. Cell Rep 35, 108978 (2021).

23. Zhang, S. et al. Genetically Defined, Syngeneic Organoid Platform for Developing Combination Therapies for Ovarian Cancer. Cancer Discov 11, 362–383 (2021).

24. Shih, I.M., Wang, Y. & Wang, T.L. The Origin of Ovarian Cancer Species and Precancerous Landscape. Am J Pathol 191, 26–39 (2021).

25. Wang, Y., Li, L., Wang, Y., Tang, S.N. & Zheng, W. Fallopian tube secretory cell expansion: a sensitive biomarker for ovarian serous carcinogenesis. Am J Transl Res 7, 2082–2090 (2015).

26. Ford, M.J. et al. Oviduct epithelial cells constitute two developmentally distinct lineages that are spatially separated along the distal-proximal axis. Cell Rep 36, 109677 (2021).

27. Lengyel, E. et al. A molecular atlas of the human postmenopausal fallopian tube and ovary from single-cell RNA and ATAC sequencing. Cell Rep 41, 111838 (2022).

28. Bartlett, T.E. et al. Epigenetic reprogramming of fallopian tube fimbriae in BRCA mutation carriers defines early ovarian cancer evolution. Nat Commun 7, 11620 (2016).

29. Ferreira, A. et al. Crucial Role of Oncogenic KRAS Mutations in Apoptosis and Autophagy Regulation: Therapeutic Implications. Cells 11 (2022).

30. Basu, D. et al. Identification, mechanism of action, and antitumor activity of a small molecule inhibitor of hippo, TGF-β, and Wnt signaling pathways. Mol Cancer Ther 13, 1457–1467 (2014).

31. Townsend, D.M. & Tew, K.D. The role of glutathione-S-transferase in anti-cancer drug resistance. Oncogene 22, 7369–7375 (2003).

32. Yan, X.D., Pan, L.Y., Yuan, Y., Lang, J.H. & Mao, N. Identification of platinum-resistance associated proteins through proteomic analysis of human ovarian cancer cells and their platinum-resistant sublines. J Proteome Res 6, 772–780 (2007).

33. Yao, F. et al. ETS2 promotes epithelial-to-mesenchymal transition in renal fibrosis by targeting JUNB transcription. Lab Invest 100, 438–453 (2020).

34. Sinh, N.D., Endo, K., Miyazawa, K. & Saitoh, M. Ets1 and ESE1 reciprocally regulate expression of ZEB1/ZEB2, dependent on ERK1/2 activity, in breast cancer cells. Cancer Sci 108, 952–960 (2017).

35. Théveneau, E., Duband, J.-L. & Altabef, M. Ets-1 confers cranial features on neural crest delamination. PLoS One 2, e1142 (2007).

36. Yi, F. et al. Opposing effects of Tcf3 and Tcf1 control Wnt stimulation of embryonic stem cell self-renewal. Nat Cell Biol 13, 762–770 (2011).

37. Guo, B. et al. The clock gene, brain and muscle Arnt-like 1, regulates adipogenesis via Wnt signaling pathway. FASEB J 26, 3453–3463 (2012).

38. Regner, M.J. et al. A multi-omic single-cell landscape of human gynecologic malignancies. Molecular Cell 81, 4924–4941.e4910 (2021).

39. Xu, J. et al. Single-Cell RNA Sequencing Reveals the Tissue Architecture in Human High-Grade Serous Ovarian Cancer. Clinical Cancer Research 28, 3590–3602 (2022).

40. Hu, Z. et al. The Repertoire of Serous Ovarian Cancer Non-genetic Heterogeneity Revealed by Single-Cell Sequencing of Normal Fallopian Tube Epithelial Cells. Cancer Cell 37, 226–242.e227 (2020).

41. Emori, M.M. & Drapkin, R. The hormonal composition of follicular fluid and its implications for ovarian cancer pathogenesis. Reprod Biol Endocrinol 12, 60 (2014).

42. Huang, H.S. et al. Mutagenic, surviving and tumorigenic effects of follicular fluid in the context of p53 loss: initiation of fimbria carcinogenesis. Carcinogenesis 36, 1419–1428 (2015).

43. Ghosh, A. et al. In Vivo Cell Fate Tracing Provides No Evidence for Mesenchymal to Epithelial Transition in Adult Fallopian Tube and Uterus. Cell Rep 31, 107631 (2020).

44. Garcia-Alonso, L. et al. Mapping the temporal and spatial dynamics of the human endometrium in vivo and in vitro. Nat Genet 53, 1698–1711 (2021).

45. Boretto, M. et al. Development of organoids from mouse and human endometrium showing endometrial epithelium physiology and long-term expandability. Development 144, 1775–1786 (2017).

46. Turco, M.Y. et al. Long-term, hormone-responsive organoid cultures of human endometrium in a chemically defined medium. Nat Cell Biol 19, 568–577 (2017).

47. Steenbeek, M.P. et al. Fallopian tube abnormalities in uterine serous carcinoma. Gynecol Oncol 158, 339–346 (2020).

48. Jarboe, E.A. et al. Coexisting intraepithelial serous carcinomas of the endometrium and fallopian tube: frequency and potential significance. Int J Gynecol Pathol 28, 308–315 (2009).

49. Kessler, M. et al. The Notch and Wnt pathways regulate stemness and differentiation in human fallopian tube organoids. Nat Commun 6, 8989 (2015).

50. Mao, Y. et al. Lentiviral Vectors Mediate Long-Term and High Efficiency Transgene Expression in HEK 293T cells. Int J Med Sci 12, 407–415 (2015).

51. Zhang, S. et al. Genetically Defined, Syngeneic Organoid Platform for Developing Combination Therapies for Ovarian Cancer. Cancer Discovery 11, 362–383 (2021).

52. Bentsen, M. et al. ATAC-seq footprinting unravels kinetics of transcription factor binding during zygotic genome activation. Nat Commun 11, 4267 (2020).

53. Ren, X. et al. Single-cell transcriptomic analysis highlights origin and pathological process of human endometrioid endometrial carcinoma. Nat Commun 13, 6300 (2022).

54. Zhang, S. et al. Both fallopian tube and ovarian surface epithelium are cells-of-origin for high-grade serous ovarian carcinoma. Nat Commun 10, 5367 (2019).

55. Goldman, M.J. et al. Visualizing and interpreting cancer genomics data via the Xena platform. Nat Biotechnol 38, 675–678 (2020).

